# Spatial mapping of pediatric brain tumors across diagnoses and relapses

**DOI:** 10.64898/2026.08.25.746606

**Authors:** Javier Escudero Morlanes, Timo-Pekka Lehto, Ludvig Larsson, Leire Alonso Galicia, Annelie Mollbrink, Alia Shamikh, Elisa Basmaci, Gabriela Prochazka, Teresita Díaz De Ståhl, Johanna Sandgren, Fulya Taylan, Bianca Tesi, Ann Nordgren, Andrew Erickson, Alastair D Lamb, Klas Blomgren, Monica Nistér, Joakim Lundeberg, Reza Mirzazadeh, Linda Kvastad

**Affiliations:** Department of Gene Technology, KTH Royal Institute of Technology, Science for Life Laboratory, Solna, Sweden; Department of Urology, University of Helsinki and Helsinki University Hospital, Helsinki, Finland; Department of Pathology, University of Helsinki and Helsinki University Hospital, Helsinki, Finland; Research Program in Systems Oncology, Faculty of Medicine, University of Helsinki, Helsinki, Finland; Department of Oncology-Pathology, Karolinska Institutet, Stockholm, Sweden; Clinical Pathology and Cancer Diagnostics, Karolinska University Hospital, Stockholm, Sweden; Department of Molecular Medicine and Surgery, Karolinska Institutet, Stockholm, Sweden; Clinical Genetics and Genomics, Karolinska University Hospital, Stockholm, Sweden; Center for Hematology and Regenerative Medicine, Department of Medicine Huddinge, Karolinska Institutet,Stockholm, Sweden; Centre for Cancer Evolution, Barts Cancer Institute, Queen Mary University of London, UK; Department of Urology, Guy’s and St Thomas’ NHS Foundation Trust, UK; Department of Women’s and Children’s Health, Karolinska Institutet, Stockholm, Sweden; Pediatric Oncology, Karolinska University Hospital, Stockholm, Sweden

## Abstract

We present a spatial transcriptomic atlas of 19 pediatric brain tumor patients spanning nine major and rare diagnoses, including seven relapses, revealing their spatial cellular and molecular organization. Each tumor section resolves into 2 - 4 recurrent spatial archetypes across 11 biological themes, with some mirroring developmental lineage patterns - for example, oligodendrocyte-lineage programs in pilocytic astrocytomas. Spatially inferred copy-number analysis identifies relapse-associated putative clones. In one rare embryonal tumor, spatial niches in the primary tumor harboring putative clones colocalized with an archetype enriched for nervous system development and glioblast-lineage programs. In one ependymoma and one pilocytic astrocytoma, relapse-associated putative clones preferentially localized to the vasculature, suggesting regrowth during relapse may be seeded by clonal selection of residual tumor cells within specialized microenvironmental niches. This resource provides an open-access spatially resolved map via an interactive viewer to inform research on pediatric brain tumor ecosystems, relapse biology, and therapeutic strategies.

## Introduction

Pediatric brain tumors (PBTs) are among the leading causes of cancer-related deaths and morbidity in children ^1^, presenting a significant clinical challenge with a broad spectrum of tumor types, a complexity in their tumor heterogeneity and recurrent nature ^2–4^. The different diagnoses exhibit diverse genomic aberrations and genetic mutations, reflecting a complex interplay between cell types, cell states, developmental processes, and the mechanisms underlying tumorigenesis ^5–13^. Recent single-cell RNA sequencing (scRNA-seq) atlases of human brain development offer a transformative lens for interpreting transcriptomic data from PBTs. One strategy is to align tumor expression profiles - either bulk RNA-seq or tumor scRNA-seq - with normal developmental cell states, to pinpoint the putative cellular origins of various tumor subtypes, track differentiation hierarchies, and reveal oncogenic programs rooted in developmental processes ^14–17^.

Another important aspect of PBTs is understanding the spatial characteristics of tumor location and tissue organization. First, the anatomical location of the tumor strongly influences clinical outcomes, with certain brain regions being more susceptible to treatment-related morbidity and functional impairment. Moreover, spatial heterogeneity within tumors - including variations in cellular composition, vascularization, and microenvironment - can affect both tumor behavior and response to therapy ^18,19^. Advanced imaging and spatial transcriptomics tools enable mapping of these variations, providing insights into tumor growth patterns, possible invasion routes, and guiding novel treatment considerations ^20,21^. This spatial context is essential for developing future therapies and strategies that maximize tumor control while preserving neurological function.

The broad and diverse biological landscape of PBTs not only complicates diagnosis and treatment but might also account for the different recurrence rates observed across pediatric brain tumors ^22^. Relapse remains a major challenge and is often associated with poor prognosis. While many primary tumors initially respond to therapy, recurrent tumors frequently display more aggressive phenotypes and resistance to standard treatments ^23^. Studying relapses provides valuable insights into tumor evolution, clonal selection, and mechanisms of resistance. Current evidence suggests that, in general, relapsed tumors may arise from minor subclones present at diagnosis or from treatment-induced selective pressures that reshape tumor biology ^19,25–27^. However, much remains unknown about the genomic, epigenetic, and microenvironmental changes that occur between primary and relapsed disease. Addressing these knowledge gaps is critical for designing more effective therapeutic strategies and improving long-term outcomes for affected children.

In this study, we included PBTs across multiple diagnoses to assess the broader utility of spatially resolved data in exploring the molecular landscape of PBTs, with a focus on dissecting their gene expression and genomic heterogeneity. We employed spatially resolved transcriptomics (SRT) using the Visium platform to generate detailed molecular profiles to understand aspects of PBT development and progression better. Our results provide new insights into the PBT tissue landscape, which are important for guiding future development of more effective, targeted, and personalized treatment strategies for this challenging and diverse group of childhood cancers.

## Results

### Molecular profiling of pediatric brain tumors

We utilized SRT technology to analyze pediatric brain tumors in 19 patients, including seven (IDs: 2, 3, 5, 9, 10, 12, 16) with both primary and relapse tumors (Fig. 1a). In total, the cohort included nine diagnoses across 13 different clinically relevant subgroups, covering low (I) and high (III-IV) tumor grades (Fig. 1b-c). Most patients were under the age of 10 at the time of sampling (Figs. 1d-e). Our dataset consists of 31 samples across 59 Visium sections, containing 62,071 barcoded spatial locations (spots) with a median gene count and median unique molecular identifier (UMI) count per spot ranging between 318-7,255 and 832-25,251, respectively. All samples were annotated by a pathologist (Supplementary Fig. 1, Supplementary Table 1-2).

**Figure 1.**
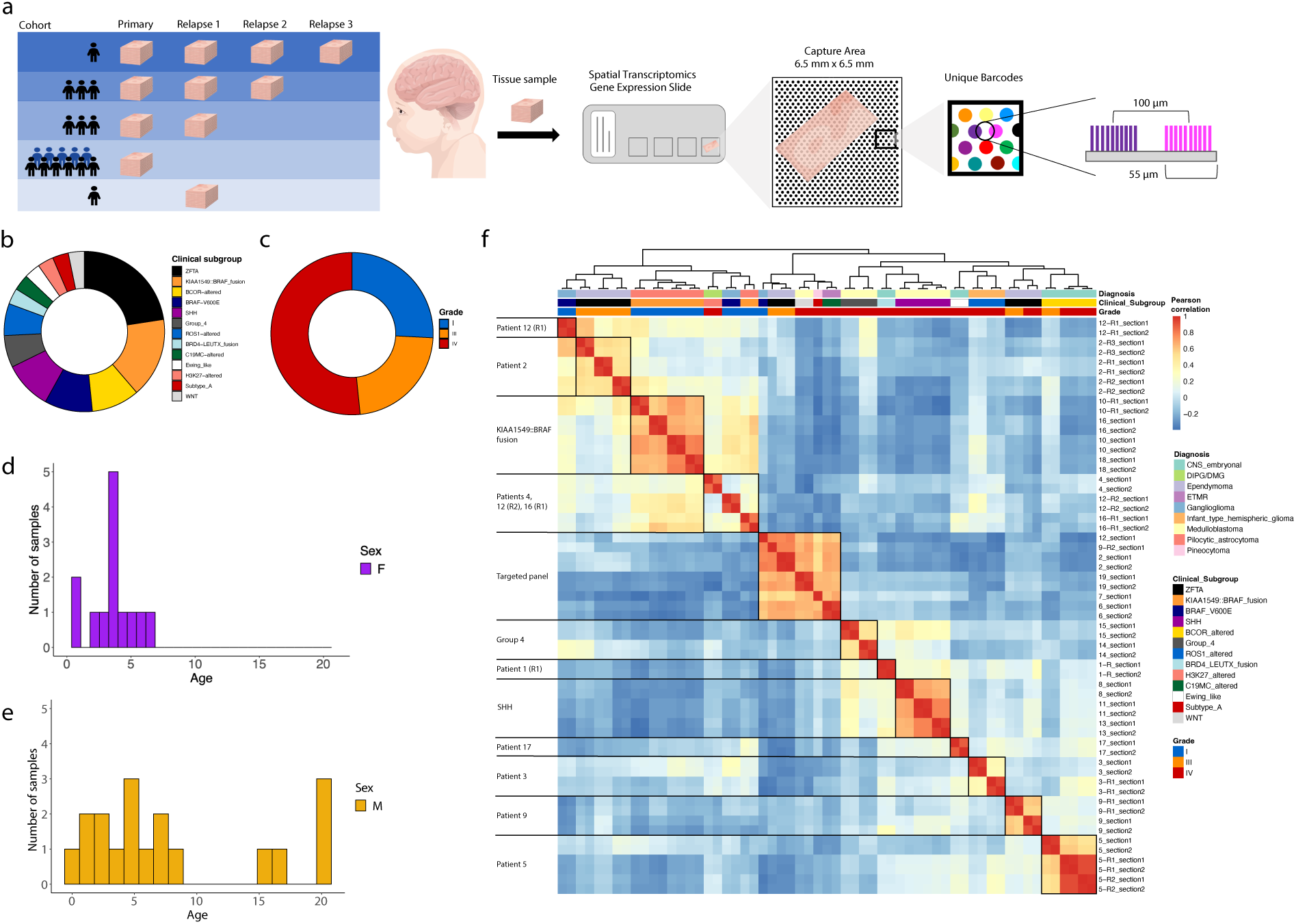
SRT on a pediatric brain tumor cohort. SRT datasets (n = 59) comprising 19 patients, and 62.071 barcoded spots. **(a)** Study design and SRT method. **(b)** Proportion of clinical subgroups across the number of samples. **(c)** Proportion of tumor grade across the number of samples. **(d-e)** Distribution of age and sex. ZFTA = Zinc Finger Translocation Associated. SHH = Sonic Hedgehog. WNT = Wingless. R = Relapse. ETMR = Embryonal Tumor with Multilayered Rosettes. DIPG/DMG = Diffuse Intrinsic Pontine Glioma / Diffuse Midline Glioma. C19MC = Chromosome 19 microRNA Cluster. **(f)** Heat map of Pearson correlation based on variable genes. Samples are named according to patient, relapse (R), and section.

For a general dataset overview, we performed principal component analysis (PCA) on pseudobulked gene expression data, focusing on the identified common variable genes. Then, we computed Pearson correlations among principal component scores (Fig. 1f). As anticipated, the strongest positive correlation occurred between adjacent tissue sections within the same specimen. We observed greater similarity between low-grade (I) and high-grade (III-IV) tumors, followed by tumor diagnosis/clinical subgroups. Notably, positive correlations were found between primary tumors and their relapse samples for patients 3, 5, 9, 10, and 16, except for the second relapse of patient 9. In contrast, positive correlations were observed only between the respective relapses for patients 2 and 12, not between the primary tumors and their relapse samples. This might be explained by a few challenging samples for polyA-based experiments (including the secondary relapse of patient 9 and the primary tumors of patients 2 and 12), which were instead processed with a transcriptome-wide targeted panel, i.e., RNA-Rescue Spatial Transcriptomics (RRST) ^28^, which excludes ribosomal and mitochondrial genes.

Finally, strong positive correlations were observed among patients belonging to three clinical subgroups: pilocytic astrocytoma with *KIAA1549::BRAF* fusion (patients, n = 3), SHH medulloblastomas (patients, n = 3), and Group 4 medulloblastomas (patients, n = 2).

### Gene expression differences between high- and low-grade tumors

The observed similarities in gene expression across tumor grades prompted further investigation. Differential gene expression analysis between grade IV and grade I tumors identified 1093 differentially expressed genes (DEGs), with 555 upregulated in grade IV tumors and 538 upregulated in grade I tumors (Fig. 2a, Supplementary Table 3). Others have previously identified similarities between transcriptomic activity of PBTs and brain developmental processes ^29^. Here, we used scRNA-seq data from the first trimester of the developing human brain ^30^ and stereoscope ^31^ to predict cell-type enhanced genes (Supplementary Table 4 and Supplementary Fig. 2), which we then overlapped with identified tumor DEGs (Fig. 2a). In grade I tumors, we found 50 genes predicted in immune-related cells and 30 genes in the oligodendrocyte lineage. For grade IV tumors, we identified 15 genes predicted to be expressed in fibroblasts and 79 genes from the neuronal lineage (Fig. 2a and Supplementary Table 5). This is in line with previous studies where low-grade tumors showed increased immune infiltration and high-grade tumors showed increased fibroblast presence ^32,33^.

**Figure 2.**
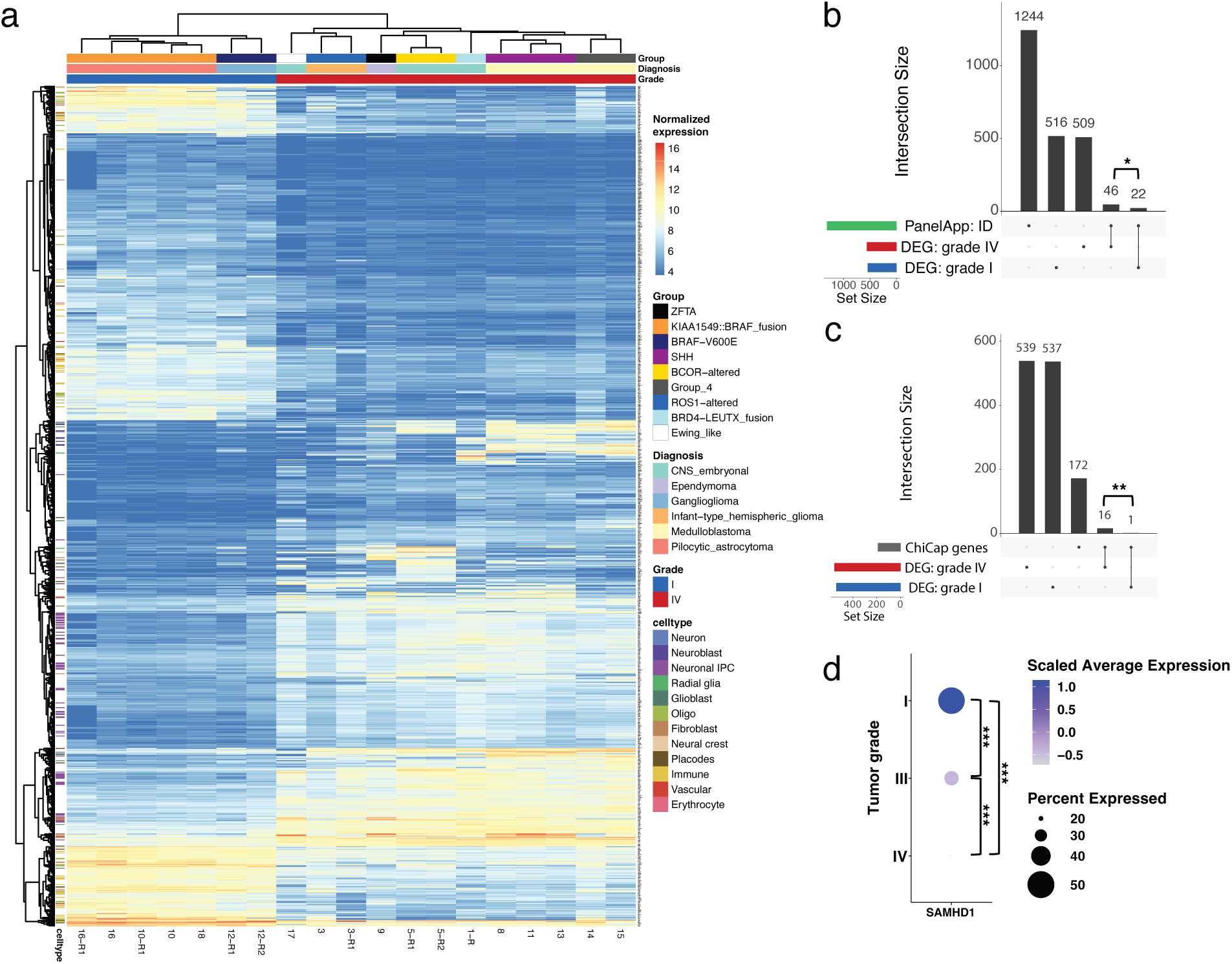
DEGs between high- and low-grade tumors across cell type enhanced and functional gene sets. **(a)** Heatmap of DEGs for high- and low-grade tumors, showing the genes predicted as most expected for each cell type from early human brain development. **(b)** DEGs compared to the PanelApp’s intellectual disability (ID) gene set. **(c)** DEGs compared to the ChiCaP’s gene list for cancer predisposition. **(d)** Gradual decrease in gene expression of *SAMHD1* across tumor grades.

It has previously been reported that children with intellectual disability have an increased cancer risk ^34^, and that germline mutations are found in at least 8.5-11% of children and adolescents with cancer ^35–37^. Thus, we examined the overlap between the DEGs from grade I and grade IV tumors with the PanelApp’s gene set for intellectual disability ^38^ (Fig. 2b, Supplementary Table 3), and the Swedish childhood cancer predisposition study ChiCaP’s gene list ^39^ (Fig. 2c, Supplementary Table 3). PanelApp divides and rates genes within their gene lists into multiple tiers with varying levels of confidence. Here we included genes rated with the highest levels of confidence ^38^. We found that DEGs from grade IV tumors, compared to grade I tumors, overlapped in significantly higher numbers with the PanelApp’s intellectual disability gene set (Fisher’s exact test, Bonferroni adjusted p-value = 0.033, odds ratio [OR] = 2.03) and with the ChiCaP’s gene list for cancer predisposition (Fisher’s exact test, Bonferroni adjusted p-value = 0.0011, OR = 15.49). We further investigated the presence of known oncogenes and tumor suppressor genes among the DEGs utilizing the OncoKB precision oncology knowledge base ^40,41^ (Supplementary Figs. 3a-d, Supplementary Table 3), finding significantly more in grade IV tumors, compared to grade I tumors, for tumor suppressor genes (Fisher’s exact test, Bonferroni adjusted p-value = 0.0022, OR = 4.65) but not oncogenes (Fisher’s exact test, Bonferroni adjusted p-value = 0.54, OR = 1.84). Furthermore, for the most broadly expressed tumor suppressor gene *SAMHD1* ^42^, we observed a significant (Pairwise Wilcoxon rank sum test, Bonferroni adjusted p-value < 2e-16) gradual decrease in gene expression across tumor grades I, III, and IV (Fig. 2d, Supplementary Fig. 3c). The specific set of tumor suppressor genes upregulated in low-grade and high-grade tumors could provide an insight into these tumors’ biological processes. We found tumor suppressor genes upregulated in low-grade tumors (e.g., *SAMHD1*, *CDKN2B, LRP1B*) that may reflect more homeostatic-like suppressor activity ^43,44^, and genes upregulated in high-grade tumors (e.g., *RAD51, POLD1, POLE, FANCA, FANCC, FANCD2, FANCG, TP53*) that may instead correspond to compensatory activity of DNA repair pathways in response to accumulating genomic damage ^45–48^. Functional enrichment analysis of the DEGs further corroborated our observations (Supplementary Fig. 3 e and f).

### Pareto task inference characterizes archetypal tumor niches

Having observed transcriptional similarities within subgroups at the pseudobulk level, we next investigated their spatial organization within tumors. Tumor growth and progression depend on the tissue microenvironment and require overcoming nutrient, cellular, and immune constraints ^49^. Using RNA-seq data, others have found that some cancer types make trade-offs between specific tasks, thereby sensitizing them to treatments targeting those same tasks ^50^. Different spatial locations, such as tumor leading edges, can create distinct environments in which different tasks can contribute to tumor cell survival ^51^. This generates a system in which tasks’ trade-offs optimize survival within a spatial niche. Characterization of these spatial niches could further our understanding of the tumor ecosystem and improve treatment outcomes.

To explore the spatial landscape across PBTs, we applied Pareto task inference (ParTI) analysis ^52^ to each section. We expected to see different niches specializing in distinct biological tasks. Since each spatially allocated spot often contains multiple cells, we also expected to sometimes find more than one type of task within the same niche. Spots from each section were fitted within a polyhedron in the PCA manifold (Supplementary Fig. 4a). Within this manifold, each vertex of the fitted polyhedron represents an archetype, which is defined as an optimal gene expression profile for a spatial biological task or niche. Consequently, proximity to a vertex implies specializing in that task and losing the ability to perform another. We found 2-4 significant archetypes for all but one section (Supplementary Fig. 4b, Supplementary Fig. 5-12 spatial archetype scores and Supplementary Table 6). In total, we found 147 archetypes across the cohort, with the majority being associated with one or more of 11 main biological themes (Fig. 3a-b). Spots closest to an archetype were used for biological characterization by i) identifying genes with the highest median difference to the remainder of the spots, ii) performing functional enrichment analysis and iii) analyzing the assigned biological processes (Supplementary Table 6). In cases where few genes were driving an archetype, or no biological function could be determined, we assigned the archetype as a panel signature. To validate archetypal themes, we performed colocalization analysis by correlating archetype gene set scores with Non-Negative Matrix Factorization (NNMF) derived factor ^53^ gene set scores using Pearson’s correlation. The most strongly correlated factors showed biological themes similar to those of the archetypes (Supplementary Fig. 13 and Supplementary Table 6). For panel signatures colocalizing with an NNMF factor of a developmental theme, we generally observed genes with known enhanced expression in neuronal and glial cells.

**Figure 3.**
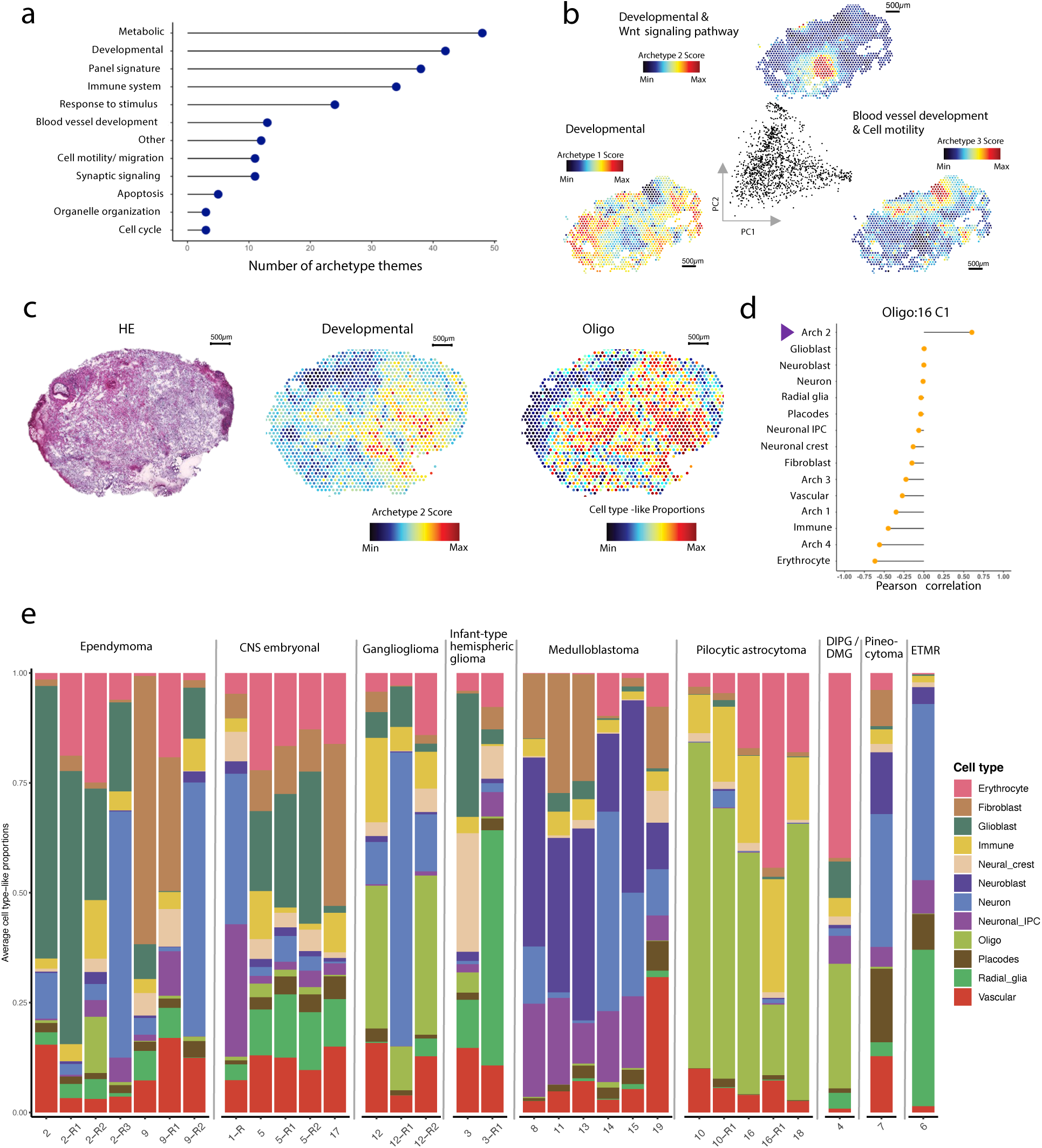
ParTI archetype analysis. **(a)** Main archetype themes across the cohort, themes occurring less than 3 times were summarized as Other. **(b)** Spatial visualization of archetype themes, and PCA manifold for patient 19 section B1. **(c)** Patient 16 section C1, tissue stained with hematoxylin and eosin, archetype 2 score, and predicted oligo cell type-like proportions. **(d)** Pearson correlations in pilocytic astrocytoma patient 16 section C1, between cell type-like proportions from the oligodendrocyte lineage and archetype scores or other cell type-like proportions. The developmental themed archetype is marked with a purple arrow. **(e)** SRT data for all samples at a pseudo-bulk level, showing inferred cell type-like proportions from the first-trimester of the developing human brain ^30^ for spots annotated as containing tumor cells.

To further validate the archetype themes, we colocalized archetypal weights with cell-type deconvolution results (Supplementary Fig. 14, Supplementary Table 7). We observed strong correlations between cell type-like proportions of the oligodendrocyte lineage and developmentally themed archetypes in DIPG/DMG (patient 4) and pilocytic astrocytoma samples (patients 10, 10-R1, 16) (Fig. 3c-d, and Supplementary Fig. 15 a-g), except in the pilocytic astrocytoma samples from patients 18 and 16-R1. Sample 18 showed high and even spatially distributed oligodendrocyte lineage–like proportions (Supplementary Fig. 16a). No developmental archetype was annotated in 16-R1, but one section’s top-correlating archetype colocalized with a developmentally themed NNMF factor (Supplementary Fig. 16b-d).

### Inferred cell type-like proportions from early human brain development identify diagnosis-specific gene expression profiles

Having observed spatial niches with gene expression patterns resembling early human brain development cell types, we further analyzed cell type–like proportions at a pseudo-bulk level, restricting to spatial spots annotated as tumor cells. Interestingly, we found that each diagnosis had a specific cell type-like proportion composition, reflecting the tumor’s underlying biological origin (Fig. 3e). During the first trimester of the developing human brain, the radial glia cells give rise to both glioblasts and neuronal intermediate progenitor cells (IPCs). The glioblasts later generate both the oligodendrocyte lineage and pre-astrocytes, whereas the neuronal IPCs differentiate into neuroblasts that later mature into neurons ^30,54^.

The medulloblastoma profiles contained large proportions of neuronal lineage-like signals, in line with previous findings, pointing towards cerebellar neurons as the tumor origin, specifically granule cell precursors and unipolar brush cells ^55–58^.

In astrocytoma profiles, including pilocytic astrocytoma and DIPG/DMG, we observed high proportions of oligodendrocyte lineage-like signals. This is in line with previous findings, which have proposed that pilocytic astrocytoma originates from the oligodendrocyte cell lineage^59^. Furthermore, we noted an interesting difference between the pilocytic astrocytoma samples and the patient diagnosed with DIPG/DMG, in which the latter showed gene expression similarities with radial glia, glioblastoma, and neuronal IPC. This could be attributed to its infiltrative nature and tendencies towards ill-defined boundaries of growth ^60^.

Ganglioglioma, a rare neoplasm containing both neuronal and glial components, has been reported to be genetically similar to pilocytic astrocytomas ^61^. Comparing cell-type-like proportion profiles between the two, we find that the ganglioglioma tumor samples exhibit similar proportions, though with increased contributions from the neuronal lineage.

In the remaining profiles, we observed increased glioblast signal in ependymomas compared to other diagnoses, and this tumor has previously been proposed to arise from an astroglial lineage ^59^. We also found that radial glia showed stronger signals in infant-type hemispheric glioma, embryonal, and ETMR tumor profiles compared to others. In pineocytoma, the highest inferred cell type-like proportions were from the neuronal lineage.

### Identifying spatially inferred copy-number variations (siCNVs) with inferCNV

Others have characterized the landscape of genomic alterations across pediatric tumors and found great interpatient variability in copy-number variation (CNV) across different brain tumor diagnoses ^62^. Furthermore, inferCNV ^63^ has been used to infer large-scale CNVs from transcriptomic data, distinguish malignant from normal cells, and to resolve tumor clonality in scRNA-seq and SRT datasets ^64–67^. Here, we apply inferCNV to identify aberrant gene expression across chromosomes as spatially inferred CNVs (siCNVs). These results were cross-referenced with available patient-matched diagnostic data of whole-genome sequencing (WGS) or whole-exome sequencing (WES). Importantly, WGS/WES and SRT were performed on distinct tumor specimens, where the relative spatial proximity of the sampled regions within the tumor is unknown. Thus, this comparison carries a caveat, as sampling different tumor regions may capture markedly distinct clonal populations ^68^.

For a majority of samples (21 of 31, 68%), biobank CNV profiles were available, with 13 samples containing CNVs and 8 samples being CNV neutral. In total, we noted 48 large scale CNVs (Supplementary Fig. 17). For siCNV analysis, randomly selected SRT spots from each tissue specimen were used to generate a cohort overview (Supplementary Fig. 18). We observed, fully or partially, 79.2% (38 of 48) of the WGS/WES derived CNVs also among cohort overview siCNVs, in addition to many non-overlapping (Supplementary Fig. 18). Non-shared CNVs between WGS/WES and siCNV could reflect noise from reference cohort selection, differences in sampling sites, and the higher spatial resolution of siCNV.

To further investigate our findings, we analyzed the medulloblastoma SHH subgroup, comparing siCNVs with WGS-derived CNVs (Fig. 4a-d). We observed 66.7% (6 of 9) of the CNVs in the siCNV data, either fully or partially. Several siCNV contained genes previously reported to be associated with the SHH subtype, such as *PIK3CA*, *PTCH1* ^69^. For patient 8 in chromosome 2, we did not observe any large siCNVs. To validate, we selected two genes (*MYCN*, *BCL11A*) from the region with the largest CNV gain, as identified by WGS data. Using TaqMan we confirmed the absence of a chromosomal gain in the tissue sample used to generate the SRT and siCNV data. This shows that diagnostic WGS/WES data from the same tumor can be informative, but differences may arise from sampling-site variation and tumor heterogeneity.

**Figure 4.**
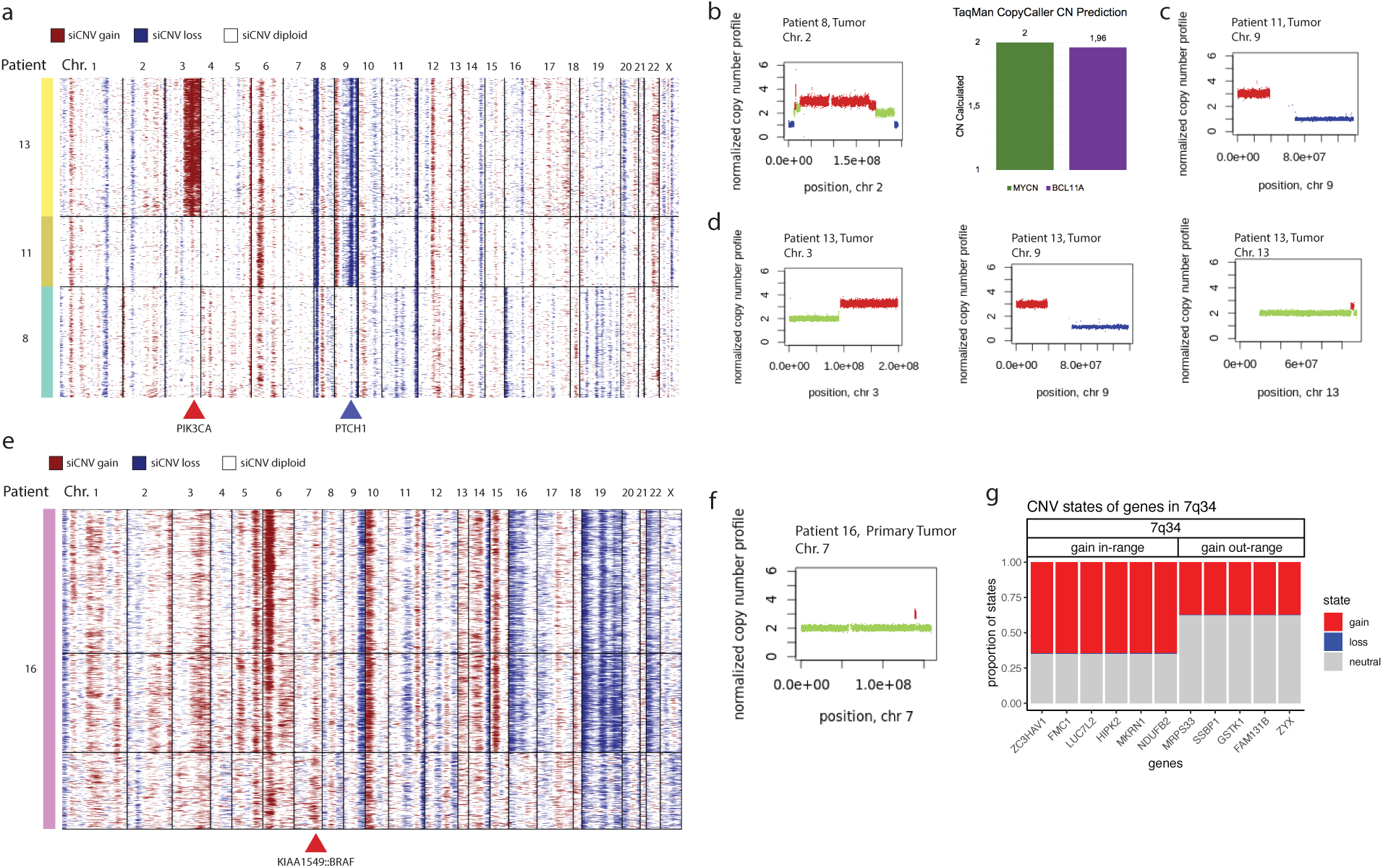
siCNV of the SHH subgroup. **(a)** siCNVs of all SHH patients, reference data from the adult dorsolateral prefrontal cortex. **(b)** Patient 8 CNV profile for chromosome 2 (left) and TaqMan CNV assay of genes from chromosome 2 (right), CN = Copy Number. **(c)** Patient 11 CNV profile for chromosome 9. **(d)** Patient 13 CNV profiles for chromosomes 3, 9, and 13. **(e)** siCNVs of patient 16, a sample diagnosed with pilocytic astrocytoma and *KIAA1549::BRAF*-fusion. Grouping of spots is performed using Leiden clustering. **(f)** Patient 16 CNV profiles for chromosome 7. **(g)** Overview of inferred HMM states across the chromosomal band where the duplication occurs in patient 16.

We also assessed the sensitivity of siCNVs by testing their ability to detect small alterations characteristic of pilocytic astrocytoma in relapse patients 10 and 16, which show the common *KIAA1549::BRAF*-fusion (Fig. 4e-g, Supplementary Fig. 17). This duplication is the driver event, where the fusion between both genes causes the constitutive activation of BRAF and a copy number gain on chromosome 7, specifically on q34 ^70–72^. WGS confirmed a small gain event on the chromosomal coordinates matching the *KIAA1549::BRAF* regions on chromosome 7 (Fig. 4f). Notably, a gain was picked up in the same region for the siCNVs (Fig. 4e-g). We next examined siCNV status across genes in 7q34, finding that genes within the KIAA1549::BRAF duplicated region were more consistently called as gained than those outside it (Fig. 4g). Thus, siCNV can capture smaller CNV patterns. Still, results should be interpreted cautiously due to potential noise.

### Spatial clonal mapping shows potential for identifying anatomical niches of relapse-associated putative clones

The development of cancer and subsequent relapses can be understood from an evolutionary perspective, in which distinct genomic and genetic aberrations give rise to different clones with varying survival capabilities. In this context, the fittest clones outgrow the rest ^73^. In the brain, others have previously identified clonal selection in, e.g., medulloblastoma relapses ^19,74^, and our SRT data provides an opportunity to explore spatial mapping of such temporal putative clonal patterns across multiple other diagnoses. In all relapse cases in this study, we confirmed from medical records (Supplementary Table 8) that tumor regrowth occurred at the same location as the previously resected tumor mass. Furthermore, having observed similarities in gene expression patterns across time points in relapse patients (Fig. 1f), we next investigated the potential that some relapses may have been driven by clonal selection of residual tumor cells regrowing at the primary site.

To identify putative siCNV temporal patterns across the relapse cases, we leveraged unsupervised phylogenetic tree reconstruction (see methods) across multiple sample time-points for each relapse patient (Fig. 5a-b, Supplementary Fig. 19). Although the phylogenetic trees suggest finer-grained clonal annotations are possible, we conservatively focused on their major structure to identify overarching temporal patterns between samples. The primary tumors were consistently found at the top of the tree, showing a clear temporal ordering. Interestingly, for all putative clones we observed various extent of spot similarity between sample time-points (Supplementary Fig. 19). This was most pronounced in a rare embryonal tumor (patient 5, Fig. 5a-b), where the primary tumor regions containing later putative clones (B and C) were most similar to relapse samples and colocalized with a developmental archetype enriched for nervous system development processes, and glioblast-like proportions (Fig. 5c, Supplementary Table 6).

**Figure 5.**
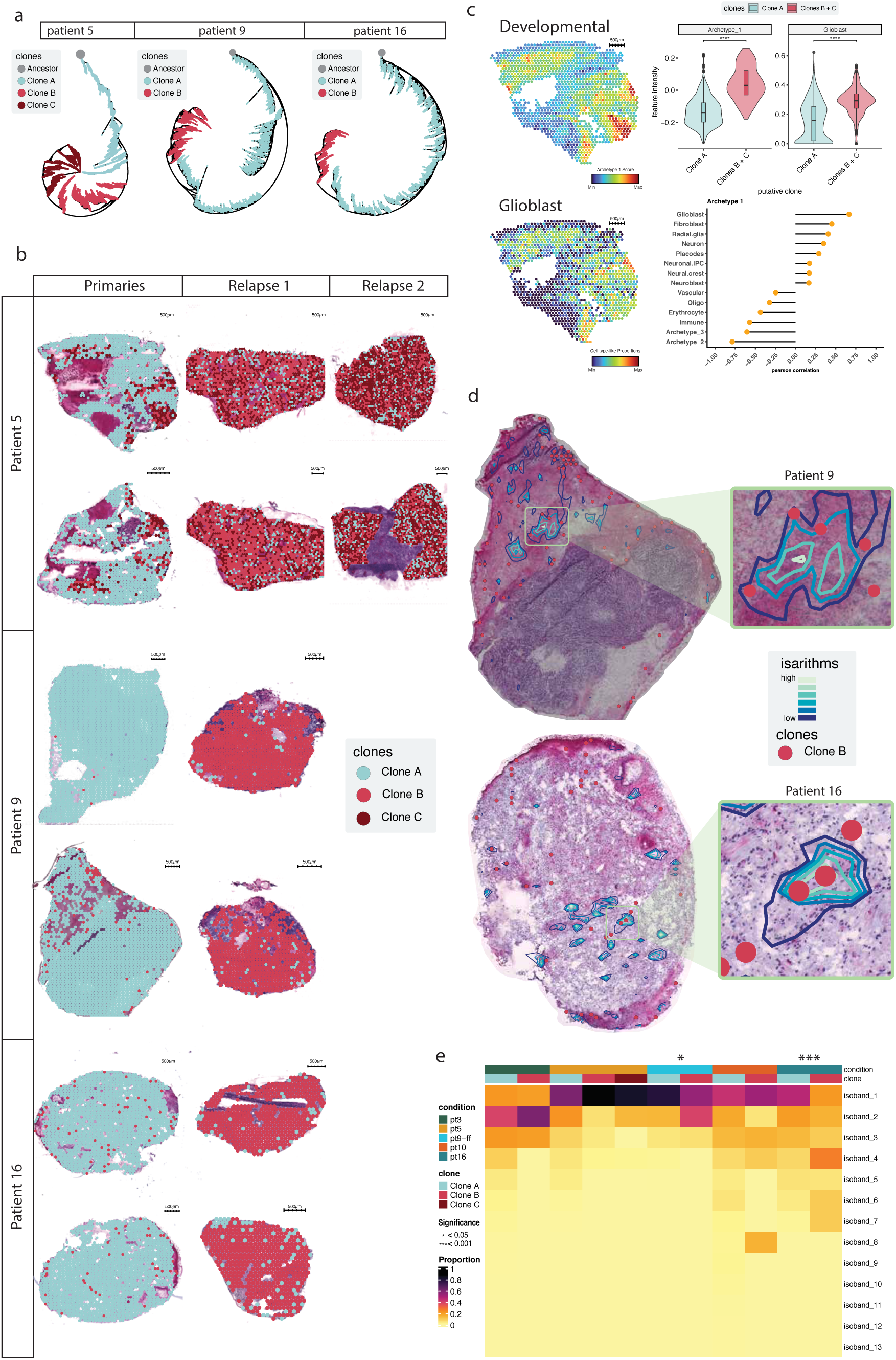
Spatial niches of tumor clones. **(a)** Unsupervised phylogenetic trees coloured by putative tumor clone assignments. **(b)** Spatial overview of putative clones. **(c)** Patient 5 section A1. Spatial visualization of selected archetype and cell type-like signals; violin plots showing contributions of archetype 1 score and glioblast cell-type like proportions across putative cancer clones (respectively: Bonferroni adjusted p-values = 5.64 x 10^-69^, 9.02 x 10^-37^); Pearson correlations between archetype 1, other identified archetypes in the section, and cell type-like proportions. **(d)** Spatial mapping of vasculature isarithms and putative late clone B in primary tumor sections of patients 9 and 16. **(e)** Proportions of spots classified in an isoband, per putative clone (for significant patients 9 and 16, respectively: permuted p-value = 0.04, 6.6 x 10^-4^, effect size = 0.092, 0.16, 95% CI = [0.02, 0.16], [0.05, 0.27]).

Given the importance of perivascular niches for tumor maintenance, dissemination, and recurrence, we investigated whether relapse-associated putative clones (clone B) in the primary tumor were associated with such niches. We developed a strategy that generates molecular isarithms from spot-level vasculature-like proportions (Fig. 5d). Isarithms define regions of equal vasculature signal, while the intervening isobands represent signal ranges. Classifying spots by isoband yielded categories reflecting both vasculature abundance and spatial proximity, with isoband 0 indicating minimal vasculature signal and the greatest distance from putative blood vessels. We next compared isoband distributions between early and late putative clones across primary tumor tissues. In patients 9 (ependymoma) and 16 (pilocytic astrocytoma), late putative clones were significantly closer to vessels than early putative clones, suggesting non-random spatial organization (Fig. 5e). Although ependymomas and pilocytic astrocytomas present relatively stable genomes ^62,75^, others have previously reported genomic and genetic differences in both types between primary and relapse samples ^76,77^. These observations suggest that a small number of residual tumor cells after resection, potentially comprising clones better adapted for survival, may seed regrowth during relapse.

## Discussion

Mapping the spatial gene expression landscape of PBTs is key to understanding their complex biology. Here, we used SRT to investigate their molecular architecture across major and rare PBT types defined by brain location, pathology, and clinical features. SRT revealed distinct tumor niches characterized by archetypal gene expression programs linked to specific biological tasks, highlighting a dynamic landscape with spatial and temporal developmental patterns.

Pilocytic astrocytoma showed colocalization between developmental archetypes and the oligodendrocyte lineage, suggesting a developmental echo. Across tumors, diagnosis-specific cell-type–like compositions reflected developmental origins, suggesting that PBTs may represent stalled or misdirected developmental programs.

Comparison of high- and low-grade tumors revealed distinct gene expression programs linked to tumor aggressiveness. Low-grade tumors were enriched for immune and oligodendrocyte lineage genes, while high-grade tumors were enriched for fibroblast and neuronal lineage genes, consistent with previous studies ^32,33^. High-grade tumors also showed greater overlap with genes associated with intellectual disability, cancer predisposition, and tumor suppression, indicating increased biological complexity.

Despite limitations in reference matching and single-region SRT sampling, siCNV analysis revealed a heterogeneous landscape of genomic alterations across tumor types, consistent with previous studies ^62^. In SHH medulloblastoma, siCNV and WGS were concordant for large CNVs. Still, we also observed differences between WGS-derived profiles from biobank sampling sites and SRT-derived siCNVs from other tumor regions, highlighting the importance of multi-region sampling.

Primary tumors and their corresponding relapses arising at the same brain location often shared a continuum in global gene expression. We observed developmentally related expression patterns between primary and relapse tumors, including similarities to cell types from early brain development. These findings suggest a degree of transcriptional persistence over time, potentially providing stable therapeutic targets, although larger cohorts are needed to validate and refine these observations. Notably, some putative relapse-associated clones were significantly closer to vessels than early clones, indicating non-random spatial organization. This finding is of particular interest given the role of perivascular niches in tumor progression and the clinical use of anti-angiogenic therapies such as Bevacizumab in low-grade gliomas, e.g., pilocytic astrocytoma ^78,79^.

Although the association between putative siCNV temporal patterns and vascular niches was modest, it supports a model in which relapse may arise through selection of residual tumor clones, potentially with enhanced survival in perivascular niches. In one ependymoma and one pilocytic astrocytoma, spatial analysis identified primary tumor regions located near vessels that corresponded to putative relapse-associated clones occupying most of the relapse-sampled tissue, highlighting perivascular niches as potential reservoirs of recurrent disease. In these tumors, relapse is often driven by remaining cells post gross total resection, and it has been reported (e.g., in ependymoma) that subsequent rounds of radio- and chemotherapy can increase mutational burden in the relapses ^77^. Future studies could assess these patterns across tumor core and tumor– normal border regions in an expanded relapse cohort. In one rare embryonal tumor, we identified primary tumor regions with relapse-like expression profiles enriched for nervous system developmental processes, suggesting that developmental and clonal trajectories may be traceable across time points. Together, these findings demonstrate the potential of spatial clonal mapping to uncover anatomical and transcriptional features of residual disease. However, interpretation is limited by the mini-bulk nature of Visium spatial transcriptomics, in which each 55 μm spot contains mixed cellular populations. Consequently, clonal assignments remain putative and may obscure finer-scale heterogeneity and true lineage relationships. These results underscore the value of higher-resolution spatial profiling, particularly at resection margins, to better understand the microenvironments that support tumor regrowth.

In conclusion, our study offers a window into the complex interplay between spatial genomics, developmental processes, and the disease biology of pediatric brain tumors. We uncovered key insights into spatial heterogeneity, potentially developmentally stalled and misdirected gene programs, and putative clonal evolution between primary and relapsed tumors. These findings may inform future research directions and guide improvements for diagnostic and therapeutic strategies.

## Methods

### Patient cohort and spatial transcriptomic datasets

Tumor tissue specimens were provided by the Swedish Childhood Tumor Biobank and stored at −80 °C until embedding for cryosectioning. The patient cohort includes 19 patients (8 female, 11 male), in total. The samples spanned all multiple tumor grades at initial diagnosis, 4 low grade (I) and 15 high grade (III-IV), 10 supratentorial (cerebrum, thalamus, tectum) and 9 infratentorial (cerebellum, pons) tumor locations, several time points (from primary and up to three relapse surgeries), ages 0.5-20.7 years, 13 different subgroups (Supplementary Table 1 and 9). In several cases, the diagnosis was updated in accordance with the WHO 2016 (PMID: 27157931) and WHO 2021 (PMID: 34185076) classifications of CNS tumors. One patient was initially diagnosed with a grade II diffuse astrocytoma in the pons, later revised to grade IV H3K27M altered DIPG/DMG. As a result, we excluded this sample from the grade I and IV DEG analysis. Spatial gene expression analysis was performed using Visium (10X Genomics), on duplicate sections from each tissue specimen (n=31). Most tissue specimens were processed with the Visium kit for fresh frozen (FF) tissues. For some specimens, due to poor RNA quality, unsuccessful library preparation with the FF protocol, or a very limited amount of tissue, the adapted RRST protocol ^28^ and the Visium kit for formalin-fixed, paraffin-embedded (FFPE) samples were used instead (Supplementary Table 1).

### Spatial transcriptomics library preparation (Visium)

The Visium Spatial Gene Expression Slide & Reagent Kit for FF and FFPE tissues (10X Genomics) was used to prepare sequencing libraries following the manufacturer’s instructions. Including the following tissue-specific steps for the FF protocol: sections were taken at 12µm and pepsin was used for 30 min. For libraries processed with the RRST protocol ^28^, samples were cryo- sectioned at 10µm, placed on Visium slides and stored at -80°C. Visium slides were removed from the -80°C freezer and placed in a thermocycler preheated to 37°C for 1-2min, followed immediately by fixation in 4% methanol-free formaldehyde (Thermo Fisher Scientific, Catalog number: 28906) solution. After fixation, Visium slides were washed twice with 1x PBS. Next, slides were air-dried, heated to 40°C for 20 min in a thermocycler, cooled to room temperature, and then stained with Hematoxylin-Eosin, optimized for each pediatric brain tissue sample. First, the tissue sections were incubated for 4 minutes with Hematoxylin, washed in MilliQ water, and air-dried. Immediately after, Bluing buffer was added and washed after 30 seconds. Then, the sections were stained with Alcoholic eosin for 30 seconds, washed in MilliQ water, and air-dried before coverslipping and imaging. Directly after imaging, slides were washed with MQ water to remove the coverslip and the glycerol, air-dried and placed inside the plastic Visium cassette. The Visium Spatial Gene Expression for FFPE kit (10x Genomics, Pleasanton, CA, USA) was used. The tissue sections were washed once with 0.1 M HCl and twice with 1x PBS. The decrosslinking step in the FFPE protocol was skipped to immediately proceed with the Probe Hybridization step following 10X Visium Spatial Gene Expression Reagent Kits for FFPE protocol for the rest of the library preparation. All finished libraries were sequenced on an Illumina NextSeq platform.

### Spatial transcriptomics data pre-processing

For libraries processed with the Visium FF protocol the following was performed. Some samples (2-R3, 10, 10-R1, 12-R1, and 12-R2) were sequenced twice, and the two corresponding fastq files were then merged into a third file. The file names of these samples are marked with v3. Using Cutadapt ^80^, read 2 was trimmed to remove both poly (A) homopolymers and the TSO adapter sequence. In brief, both the partial and full-length TSO adapter sequences were removed by setting a non-internal 5’ adapter (error tolerance = 0.1 and minimum overlap = 5) to the TSO sequence (AAGCAGTGGTATCAACGCAGAGTACATGGG). By setting a sequence of 10 A’s for a 3’ adaptor (minimum overlap = 5) the polyA homopolymer sequences were removed. Then, the trimmed read 2 and the raw read 1 fastq files were run through the Spaceranger pipeline (v1.0.0, 10X Genomics) and reads were mapped to the human reference genome (GRCh38, release 93). Visium sample V10S29-107_D1 required manual alignment of bright field image. Samples were filtered using the *InputFromTable* function in STUtility ^81^(v0.1.0), all spots containing less than 500 UMI counts were removed and genes were removed if they were present in less than 5 spots or had a total UMI count below 100.

### Correlation heat map of variable genes across sample cohort

Analyses were carried out using the STUtility (v0.1.0) and Seurat (v3.2.2) R packages ^81,82^. To identify variable genes, samples were normalized using a variance-stabilizing transformation implemented in the *SCTransform* function from Seurat. Once the top 3000 variable genes had been identified, the expression profiles of these genes for each Visium sample were first aggregated into “bulk” expression vectors by summarizing the total UMI counts for each gene. Then, library size normalization was performed, where each such “bulk” expression vector was normalized to have the same total count according to: 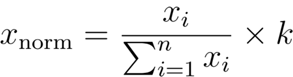, where ‘i’ is a gene, ‘*n*’ is the number of genes, and ‘*k*’ is the average of the summarized “bulk” expression vectors. After normalization, the “bulk” expression vectors were logarithmized (*log1p*), dimensionality reduction was implemented using principal component analysis with the *prcomp* function from stats, Pearson correlations were calculated and plotted as a heat map.

### Inferring cell type proportions in SRT data

Celltype proportions were inferred using stereoscope (v0.3.1) ^83^. Processed scRNA-seq data was retrieved from the Human Developmental Brain Atlas ^30^. The data is stored in a file named “*HumanFetalBrainPool.h5*”, which contains, amongst other features, cell annotations and the expression matrix. A random subset of 870 cells per annotated cell type was selected to reduce the dataset size. From this subset, genes with fewer than 50 total counts across all cells were removed. The resulting count matrix was processed using Seurat v4.3.0. Log normalization of the scRNA-seq data was performed using *NormalizeData()* with a scalefactor of 10000. The 5000 most variable genes were computed using *FindVariableFeatures(selection.method = “vst”, nfeatures = 5000),* and exported to for processing with stereoscope. Data was further reduced by randomly selecting 250 cells from each cell type to define the training data for deconvolution.

To prepare the Visium dataset for deconvolution (query data), we excluded all spots with a total UMI count below 500, as well as genes that had less than 100 total UMIs and were present in less than 5 spots. This filtering was done with STutility’s *InputFromTable()* with the following parameters: *minUMICountsPerGene = 100, minSpotsPerGene = 5, minUMICountsPerSpot = 500*. Stereoscope was run on a NVIDIA A100-SXM4-80GB, with 30000 epochs both for training and deconvolution.

### Identifying genes relevant for a celltype

As part of the standard output of stereoscope, a R*.tsv file that stores the rates for each gene in each celltype, and a logits*.tsv file are given. To define the genes most relevant to each cell type identified by stereoscope, we computed the expected value for all genes in each cell type. Given that in the negative binomial distribution, which is the model assumed by Stereoscope for gene expression data, the mean is defined by *mean = r(1-p)/p* and the general definition for a logit is *logit = log(1-p)/p*, the expected (mean) expression of a gene in a cell type is *mean_exp = rate*exp(logits)*. Based on this expected expression, the 400 most expected genes for each cell type were selected.

### Differential gene expression analysis between grade I and grade IV tumors

Differential Expression Analysis (DEA) was done between Fresh Frozen samples annotated as grade IV and grade I, using the R library DESeq2 (v1.30.1)^84^. Initial filtering was done by keeping exclusively those spots annotated by the pathologist as “tumor”. The spatial gene expression matrix for each sample was collapsed spotwise, generating pseudo-bulk count matrices that contain the total raw counts for each gene across the tissue section. Mitochondrial and ribosomal protein coding genes, as well as genes with less than 10 total counts across all samples were filtered out.

DEA was performed, with grade I tumors serving as the reference. Since in our dataset there are 2 tissue sections per tumor sample, we considered those as biological replicates, which we collapsed. Differentially expressed genes with a log-fold change equal or greater than 1 and with an adjusted p-value equal or lower than 0.05 were kept. Visualization of DEA results was performed using the *pheatmap()* function from the pheatmap R library (v1.0.12)^85^. As input for the plotting function, section-wise scaled gene counts done with DESeq2’s *vst()* were provided. Genes were sorted by decreasing log-fold change and annotated according to the cell type for which the expected expression was the greatest, as outlined in the section above.

### Comparing differentially expressed genes with intellectual disability and cancer predisposition gene sets

To analyze the relationship between the categorical values of DEGs from High/Low grade tumors and their overlap with the Genomics England (GEL) panel from PanelApp’s intellectual disability (ID) gene set ^38^(v4.4), the ChiCaP gene list for cancer predisposition ^39^ and the OncoKB cancer gene list ^40,41^, we employed Fisher’s exact test using *fisher.test* (R version 4.5.0), results were rounded of to two decimals. PanelApp rates genes according to their GEL status, which is divided into three categories based on evidence supporting their association with a specific phenotype or disease. Here, we selected to only include the ID GEL 3 category (highest level of confidence). We computed in total four tests for the following gene lists and their relationship to the number of DEGs from High/Low grade tumors; i) the ChiCaP genes, ii) the ID GEL 3 category genes, iii) the OncoKB tumor suppressor genes, and iv) the OncoKB oncogenes. We used Bonferroni to correct for multiple testing based on the number of tests.

### Pareto task inference with ParTI archetype analysis

We used the ParTI MATLAB software package ^52^ for the archetype analysis. Briefly, data for each SRT section was log normalized using the *NormalizeData* function from Seurat. The positions of the archetypes (vertices) were determined by ParTI in PCA space, computed on each section’s gene expression data. These archetypes form a polyhedron, a geometrical figure with flat sides and sharp vertices, which contains the data points. How well the polyhedron fits the data PCA manifold is computed with a *t*-ration test ^52,86^ (which compares the fitted data to a randomized dataset) to assess statistical significance (p < 0.05). For characterization of the archetypes with ParTI we used two continuous features: i) the expression of MSigDB pathways (h.all.v7.5.1.symbols.gmt and c5.all.v7.5.1.symbols.gmt) containing at least 10 genes, and ii) the expression of individual genes within each dataset. ParTI was run using the default Sisal algorithm. We tested if these features values were significantly elevated in spots close to the archetypes (Mann–Whitney U test, two-sided), using enrichment bin sizes of 20-50 closest spots (Supplementary Table 6). The false discovery rate (FDR) was set to FDR < 10% using the Benjamini-Hochberg procedure.

### Gene set scoring, colocalization and functional enrichment analysis of ParTI archetypes and NNMF factor genes

All gene set scores were calculated using the *AddModuleScore* function from Seurat on the top 200 genes contributing to each archetype or factor. Genes from the ParTI analysis were first sorted based on their median difference (from highest to lowest value), then the top 200 significant genes were used (or all genes if fewer than 200 significant total genes). Colocalization analysis was performed by calculating Pearson correlations; high positive correlations were interpreted as colocalization and were visually verified by plotting the respective scores back on top of the tissue sections. The functional enrichment analysis was performed with g:Profiler2 (v0.2.0, Ensembl 106, Ensembl Genomes 53)^87^, using the *gost()* function with an ordered query, the accompanying g:SCS method for multiple testing correction, and an adjusted p-value threshold of 0.05. Results were sorted by smallest p-value, and the top 20 biological processes were used to characterize the archetype and factor themes.

### Inference of chromosomal copy-number variants

Data was analyzed using the inferCNV (v1.18.1) R package ^63,88^. Data pre-filtering was performed using the STUtility (v0.1.0) and Seurat (v3.2.2) R packages. QC of ST data was done depending on the analysis at hand.

### Establishment of a normal tissue reference

A crucial step of running inferCNV is establishing a reference that matches the data in which CNVs wish to be computed. Our cohort is both rare and precious, making it so that no publicly available SRT pediatric brain samples are available, and so that no extra healthy tissue can be sectioned off during surgery. Therefore, a reference set for the inferCNV analysis was constructed by randomly selecting 250 spots per section from a pool of Visium spots comprising 12 sections from the adult human dorsolateral prefrontal cortex ^89^. This reference was used for all subsequent inferCNV runs.

### InferCNV analysis of a cohort overview and SHH subgroup

Input objects were created using inferCNV’s *CreateInfercnvObject()* function, excluding the mitochondrial genome. Analysis was performed by invoking inferCNV’s *run()* function with the following arguments (unless specified, arguments took default values): *cutoff = 0.1*, *cluster_by_groups = TRUE*, *HMM = TRUE*. For the cohort overview we randomly selected 360 spots from each tumor tissue specimen and manually ordered the data for the final inferCNV output by patient and clinical subgroup. Comparison with diagnostic WGS/WES profiles excluded the Y chromosome due to sparse inferCNV signals. For the SHH diagnosis subgroup, we only removed spots that i) did not include tumor cells, as annotated by a pathologist, and ii) were in locations where the tissue was folded; all others were included in the analysis.

### Spatiotemporal perspective of CNVs across relapses

To study the potential evolutionary relationship between primary and relapse tumors of the same patient, inferCNV was run on sections from selected patients (2, 3, 5, 9, 10, 12, 16 - Supplementary Table 9) of matching technologies to minimize technical noise. For example, in patient 2 two different inferCNV runs were done: one with the primary tumors (processed via the FF protocol), and another one with the relapse tumors (processed via RRST). Visium data was curated by removing spots that i) were not annotated as *tumor* by the pathologist, ii) where tissue folding happened, and iii) had less than 500 total UMI counts. Genes satisfying the following criteria were kept: common between reference and query datasets; more than 0 total counts in each individual section; more than 100 UMI counts, and present in at least 5 spots across the cohort.

Once the query and reference datasets were set up, inferCNV objects for each selected patient were created, excluding the mitochondrial chromosome, with *CreateInfercnvObject()*. To generate a file containing genomic coordinates, the functions from the R package biomaRt (v2.58.2) *useEnsembl()* and *getBM()* were used ^90,91^.

With the aim to generate the most detailed report of the tentative CNV status of each gene in each spot, while compromising for computational feasibility, the *run()* function was called with the following arguments (unless specified, arguments took default values): *cutoff = 0.1, cluster_by_groups = FALSE, HMM=TRUE, HMM_type = “i6”, HMM_report_by = “cell”, analysis_mode = “subclusters”, tumor_subcluster_partition_method = “leiden”, leiden_resolution = res, useRaster = FALSE, plot_probabilities = FALSE, num_threads = 100.* Transition probabilities were computed at the subcluster level due to computational limitations in computing them at the spot level. The resolution of Leiden clustering was determined empirically as a compromise between clustering resolution and inferCNV runtime, with the following resolutions decided for each run: patient 2 primaries, 0.005; patient 2 relapses, 0.002; patient 3, 0.004; patient 5, 0.0008; patient 9 FF sections, 0.0006; patient 9 RRST sections, 0.03; patient 10, 0.008; patient 12 primaries, 0.05; patient 12 relapses; 0.005; and patient 16, 0.0007 (Supplementary Fig. 20).

### CNV analysis of WGS/WES data

The WGS/WES data were obtained from the Swedish Childhood Tumor Bank. The WGS libraries were prepared using Illumina TruSeq PCR-free reagents. Samples were sequenced using 2 × 150- bp paired-end reads, on a HiSeqX v2.5 or NovaSeq 6000 instrument (Illumina). The WES libraries were prepared using the Agilent SureSelect Human AllExonV5 protocol (Agilent), and sequenced using 2×100 bp paired-end, on a HiSeq 2500 instrument (Illumina). The DNA sequencing data were then processed with Sarek, using the GATK best-practice recommendations ^92^, performed on UPPMAX Clusters at Uppsala University. In brief, FastQC (https://www.bioinformatics.babraham.ac.uk/projects/fastqc/) was used for quality control of the FASTQ files, the human reference genome sequence (GRCh38/hg38) was used for alignment of short reads, using bwa-mem with the ALT-aware option turned on ^93^, GATK tools (https://github.com/broadinstitute/gatk) were used for sorting of reads, base quality score recalibration, joint realignment of reads around indels, and marking of PCR duplicates with GATK MarkDuplicates. Control-FREEC ^94^ was used to generate tumor CNV profiles. Matched normal samples from peripheral blood mononuclear cells were used to call somatic CNVs.

### TaqMan assay

DNA was extracted from five sections of an OCT block from patient 8 using a Qiagen DNA extraction kit. For copy number analysis (CNA), we employed two distinct TaqMan probes targeting the MYCN and BCL11A genes on chromosome 2. These, along with a reference probe for RNaseP, were purchased from Thermo Fisher Scientific. Specifically, the MYCN probe used was the TaqMan CNV Assay (Hs07534295_cn), the BCL11A probe was the TaqMan CNV Assay (Hs00201049_cn), and the reference probe was the TaqMan Copy Number Reference Assay (4403326). The experimental procedure, including the analysis, adhered to the standard TaqMan protocol provided by Thermo Fisher Scientific, with no significant deviations from the established protocol during the experiment.

### Cross-reference of iCNV predictions and WGS results

HMM states were queried for overlap with WGS results for patients 10 and 16, both diagnosed with pilocytic astrocytoma, which showed a KIAA1549::BRAF gene fusion via WGS. HMM states and genes chromosomal positions were contextualized using biomaRt’s *useEnsembl()* and *getBM()*.

The gene-states were filtered to keep only genes in chromosome 7 arm q band 34 (7q34), which is the fusion region. The specific region affected by the fusion was identified by harmonization of WGS and inferCNV results. Since HMM states are reported only at the gene level, there is a mismatch between the coordinates from WGS and those from inferCNV. Thus, this region was characterised by computing the closest inferCNV start to that one reported by WGS; and likewise for the end.

Next, HMM states were reclassified in 3 possible labels: “loss” when the state was lower than 3, “neutral” when identical to 3, and “gain” as greater than 3. This reclassification is a simplified version of inferCNV’s internal classification of the different HMM states that can be reported. The number of labels per gene across the cohort of spots was computed, and genes were ordered along the x-axis by increasing chromosomal start position. In patient 16, genes KIAA1549 and BRAF did not meet the QC cutoff set by the inferCNV *run()* function due to very low expression levels, so CNVs were not computed for either gene. Thus, the effects of the gene fusion are explored by querying the HMM states of all genes between KIAA1549 and BRAF and comparing them with those of all remaining genes in 7q34.

### Construction of phylogenetic trees across relapse cases

Phylogenies were reconstructed from the inferCNV data using maximum parsimony ratchet from the R package phangorn (v2.11.1), specifically functions *pratchet()* and *acctran()* ^95,96^. The input used for building the trees were the HMM states coming from step 19 of inferCNV, specifically the files named 1*9_HMM_pred.Bayes_NetHMMi6.leiden.hmm_mode- subclusters.Pnorm_0.5.infercnv_obj*. Tree objects were formatted into a dataframe using a custom wrapping function that takes the phylo objects given by phangorn and converts them into dataframes with non-overlapping x and y coordinates for each node. A synthetic Ancestor node is also added at the top of the tree.

### Clonal classification

Tumor clones were defined via visual inspection of the phylogenetic trees’ morphology, aided by metadata that indicated the tissue section of origin of any spot. A maximum granularity of three levels (interpreted as putative tumor clones) was established: early/A, late-middle/B and advanced/C. This was done by identifying the terminal nodes of a given clone and gathering all downstream nodes connected to it. Section-specific enrichment of the identified clones was initially explored via pie charts (Supplementary Fig. 21).

### Clonal enrichment of transcriptomic programs

To assess if archetypal weights and cell type-like proportions showed specific enrichment for putative clones in replicate 1 of the primary tumor sections of patient 5, the R package *rstatix* was used (v0.7.3) ^97^. One-sided Wilcoxon test was run using the *wilcox_test()* function with *alternative = “less”*. Reported p-values were adjusted with Bonferroni correction invoking *adjust_pvalue()*.

### Geographical analysis

To classify vascular cell-type-like transcriptomic signatures into spatial regions, contour plots were explored. Due to the honeycomb grid pattern on the Visium slides, contour plots cannot be computed directly. Thus, it was first necessary to transform the honeycomb grid into a regular grid, which was achieved by spatially interpolating the vascular cell-type-like signature.

### Spatial interpolation

Given the structure of the Visium array, one can conceptualize that spatial interpolation will “fill in the gaps” between spots to generate a full, continuous spatial array. Spatial interpolation was performed on the relative array coordinates of each Visium spot, rather than on the real image coordinates. This was decided because small rotations were present in the pixel coordinates of Visium samples, which could be avoided by using the relative array coordinates provided by spaceranger.

We generated a regular grid from the Visium data on top of which contour plots could be computed by using akima’s (v0.6.3.6) *interp()* function ^98^. Since the aim was to “fill in the gaps” between spots in a Visium array, and considering that the relative array coordinates are used, the minimal distance at which to space the interpolated grid was set to 1. The akima’s *interp()* function was invoked with the following arguments: *x = x* is a vector storing the spatial x coordinates of the original Visium data; *y = y* is a vector storing the spatial y coordinates of the original Visium data; *z = z* is a vector storing the proportions of vascular celltype-like signal; *xo = x_seq* is a vector storing the x coordinates of the regular grid that will be returned as an output; *yo = y_seq* is a vector storing the y coordinates of the regular grid that will be returned as an output; *linear = TRUE* indicates that linear interpolation is used; *duplicate = mean* means that if duplicate spatial coordinates are found, the method will return the mean of the feature value in the duplicates. *xo* and *yo* are vectors defined by “filling in the gaps” between spots in the original array coordinates. Spots outside of the convex hull of the tissue section were removed post-interpolation.

### Considerations regarding isarithms and isobands

Isarithms are the lines in contour plots, which represent a 3D surface by connecting points with equal feature values. The term “isarithm” is a generalization of these lines, which are usually named depending on the feature value of the plot. In our case, the isarithms represent points of our tissue section that have constant values of vascular cell-type-like signature. Isobands, on the other hand, are the filled areas between contour lines. A key difference between isarithm and isoband is that the isarithm will represent a specific value, whereas the isoband is defined by the range of values between the two isarithms that conform it. Knowing this, the interpolated spatial data will be used to define isarithms, and Visium spots will be classified in isobands by utilising the values set in the isarithms.

To ensure that the computed isobands reflect the same values across the several tissue sections in each of our cohorts, contour bands were computed simultaneously across all Visium sections of a specific patient included in the inferCNV run (Supplementary Table 9). Otherwise, the same isoband label can differ in the feature value that it expresses across samples. This behaviour is expected, as neither the exact values or spatial distribution of the vascular cell-type-like has to be identical across multiple tissue sections. A Common Coordinate Framework (CCF) for the sections of interest was defined, consisting of offsetting the relative array coordinates of each tissue section by 100 units in the cartesian x or y axes. When computing isarithms in this CCF, contour bands remain section-specific but are harmonized across all sections. Classifying spots by isoband yielded categories reflecting both vasculature abundance and proximity, with isoband 0 indicating minimal vasculature signal and greatest distance from putative blood vessels.

### Spatial categorization of celltype-like proportions

Isobands were generated on the interpolated vasculature celltype-like proportions using ggplot2’s *geom_contour_filled()* (v3.5.2). Their ranges were extracted from the plot using *layer_data()*, and applied to categorize the celltype-like proportions of the original (non-interpolated) Visium data using the *cut()* function with *right = TRUE*. As a result, each Visium spot is assigned a specific isoband. If the feature value of a Visium spot is lower than the lower bound of the isobands, the spot is assigned to the lowest isoband. Conversely, if a feature value is greater than the greater end of the isobands, it gets assigned to the greatest isoband. The greater the value of an isoband, the stronger the vasculature signature is. Simultaneously, the lower the value of the isoband is, the further away from vasculature a spot is. Spatial visualization of isarithms on top of the tissue image was done by overlapping the pixel coordinates of the contour lines on plots generated using semla’s *MapLabels()* (v1.3.1) ^99^.

### Spatial enrichment of cancer clones around perivascular niches

To assess whether the identified tumor clones showed specific localisation on/around perivascular regions in primary tumor sections, the relationship between tumor clones and vasculature isobands was explored. Specifically, this meant testing on a per-patient basis whether spots in primary tumor sections identified as relapse-associated clones showed increased isoband classifications. To that end, within primary tumor samples, vasculature isobands classified as ‘isoband_0’ were excluded, as they indicate absence of vasculature cell type-like signature.

Permutation tests were run using coin’s *independence_test()*, with arguments *distribution = approximate(nresample = 100000L)* and *alternative = “greater”* ^100^. Effect size was computed using rstatix’s *wilcox_effsize()* with *paired = FALSE* and *alternative = “greater”* (v0.7.3). Results were visualized using the function *Heatmap()* from the R package ComplexHeatmaps (v2.15.4)^101^.

## Code availability

For trimming of read 2 to remove TSO and polyA homopolymers: https://github.com/ludvigla/VisiumTrim/blob/main/TSO_polyA_trimming.sh.

Code required to generate all figures in the manuscript can be found in https://github.com/jemorlanes/PBTs_code. A permanent version of the code is stored at KTH’s Data Repository (Data availability).

## Data availability

Raw or analyzed sequencing data files from the childhood brain tumour samples can be made available for authorized users after an approved formal application, including relevant ethical permits, to the Swedish Childhood Tumor Bank (Barntumörbanken), in line with GDPR regulations. All other additional data and material e.g., spot-level pathology annotation metadata, ParTI archetype output files, count matrices, high resolution histological images, space ranger output files and a permanent version of the code, are available in the KTH Data Repository (https://datarepository.kth.se/records/stbwn-0px47). The Visium spatial transcriptomics datasets can be interactively explored through https://pbt.serve.scilifelab.se/, with specific instructions for navigation found in the provided guideline.

## Additional information

### Ethics statement

This study was approved by the Regional Ethical Review Board (EPN), Stockholm, Sweden (DNR 2018/3-31 and 2022-03649-02, Monica Nistér). Tissue specimens were obtained from the Swedish Childhood Tumor Bank, where samples from human subjects had been collected with written informed consent for research purposes.

### Author contribution

LK, RM, MN and JL initiated the project. LK participated in study design, performed data analysis, assisted with pathological analysis, helped to procure ethical approval and funding and wrote the manuscript. RM participated in study design, performed experiments and wrote the manuscript. JEM developed and performed data analysis and wrote the manuscript. LL provided bioinformatic advice and early data analysis. LAG and AM performed experiments. AS, KB, and MN contributed with pathological analysis and biological insights. MN also helped to procure ethical approval and funding. EB, GP and JS contributed with biological insights and tissue provision. TDDS contributed with computational analysis. JS and TDDS also contributed with access to genomic data. FT, BT and AN contributed with biological expertise. TPL, AE and ADL provided guidance and support establishing the iCNV analysis. JL contributed with study design, writing of the manuscript and procured funding. LK and RM provided project guidance and supervision.

## Acknowledgments

The authors acknowledge The Swedish Childhood Tumor Biobank, supported by The Swedish Childhood Cancer Fund, for access and handling of patient biobank material and sequencing data, the National Genomics Infrastructure (NGI), Sweden for providing infrastructure support. This work was supported by the National Bioinformatics Infrastructure Sweden (NBIS) at SciLifeLab. NBIS is a national research infrastructure funded by the Swedish Research Council (Vetenskapsrådet Dnr 2023-00153), SciLifeLab and others. Furthermore, we would like to thank the following funding agencies: The Swedish Childhood Cancer Fund, The Swedish Research Council, The Swedish Cancer Society, Swedish Foundation for Strategic Research, The Erling Persson Foundation, Knut and Alice Wallenberg Foundation and European Research Council Advanced Grant. We thank Mattias Karlen for artistic contributions.

## Supplementary Tables

**Supplementary Table 1. Patient metadata.** Summary of metadata for the 19 pediatric brain tumor patients.

**Supplementary Table 2. SRT dataset QC data.** Summary of QC data from the 59 SRT datasets. Two tissue sections were taken from each tumor sample. N/A = library preparation and/or sequencing not successful.

**Supplementary Table 3. DEGs between low- and high-grade tumors.** The pseudo-bulk 1093 DEGs between the grade I and grade IV tumor samples of the patient cohort, and their gene list overlap with PanelApp’s gene set for intellectual disability ^38^, the Swedish cancer predisposition study ChiCaP’s gene list ^39^, known oncogenes and tumor suppressor genes among OncoKB precision oncology knowledge base ^40,41^.

**Supplementary Table 4. Stereoscope cell type enhanced genes.** The top 400 expected genes per cell type, predicted by stereoscope ^83^, including annotations for most expected gene per cell type based on gene expression from the first-trimester of the developing human brain ^30^.

**Supplementary Table 5. Tumor DEGs overlap with early brain development cell type enhanced genes.** The predicted enhanced cell type genes from the first-trimester of the developing human brain ^30^, which also overlapped with the DEGs between low- and high-grade tumors in Figure 2a.

**Supplementary Table 6. Pareto task inference analysis.** Analysis and biological characterization of the archetypes in the pediatric brain tumor cohort.

**Supplementary Table 7. Stereoscope predicted cell type-like proportions.** Metadata for SRT spot level annotation across the pediatric brain tumor cohort.

**Supplementary Table 8. Patient medical metadata.** Patient sample level information for treatment, survival, and relapse status.

**Supplementary Table 9. Tissue sections per relapse-focused inferCNV analysis.** Which samples were put together for the identification of shared siCNV profiles across tumor relapses.

