## Supplementary figures and images for "Spatial mapping of pediatric brain tumors across diagnoses and relapses"

### Supplementary Figure 1

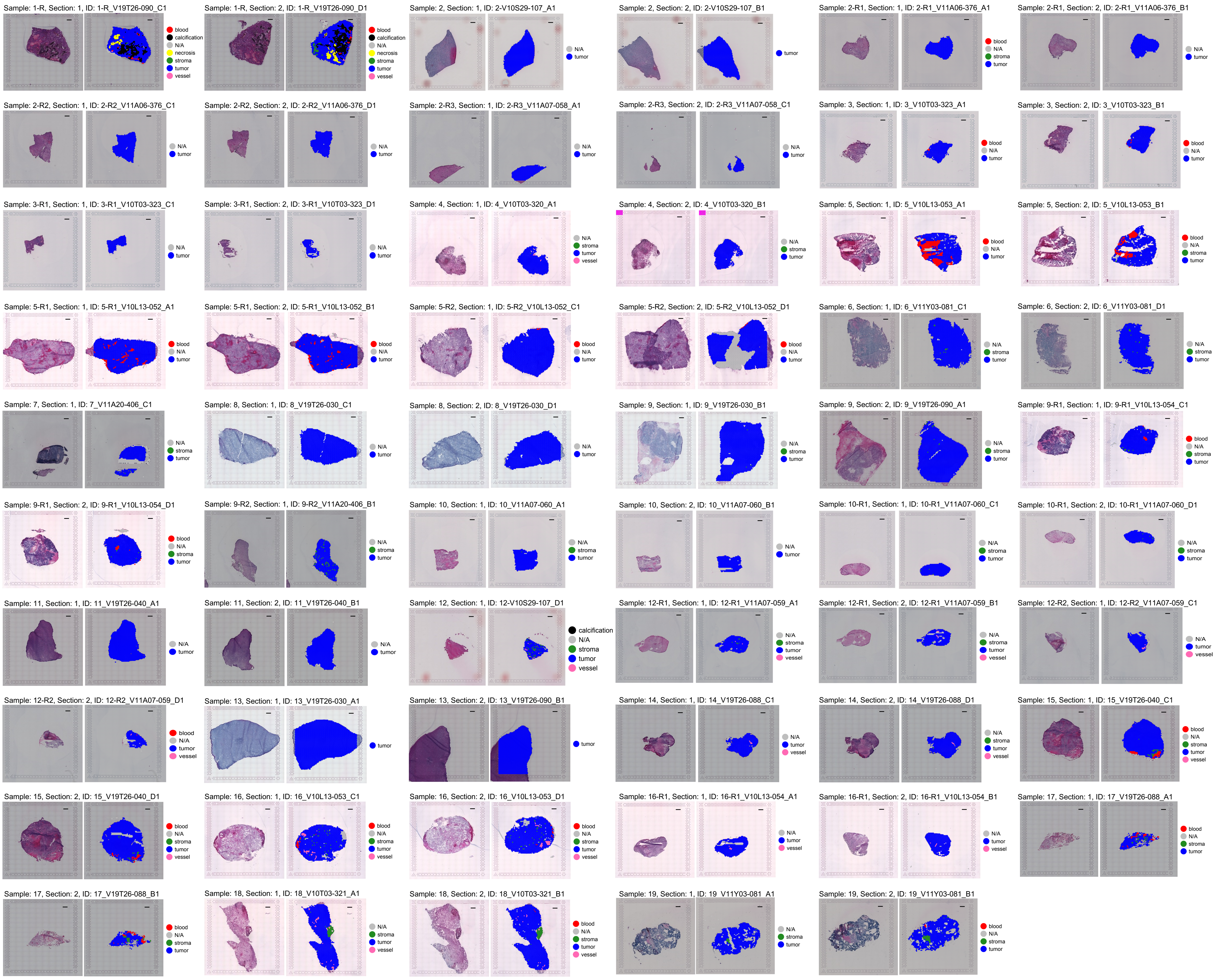

### Supplementary Figure 2

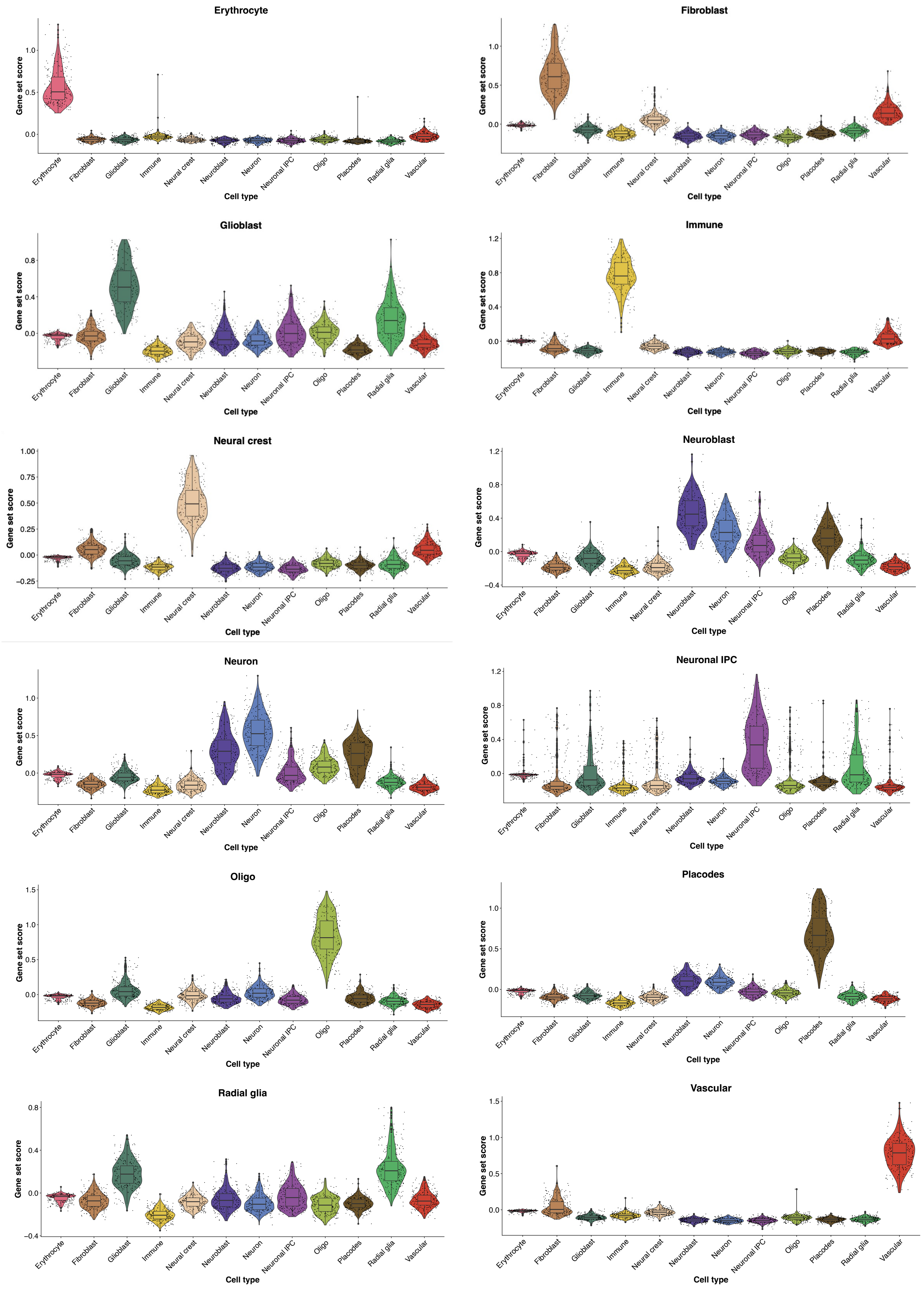

### Supplementary Figure 3

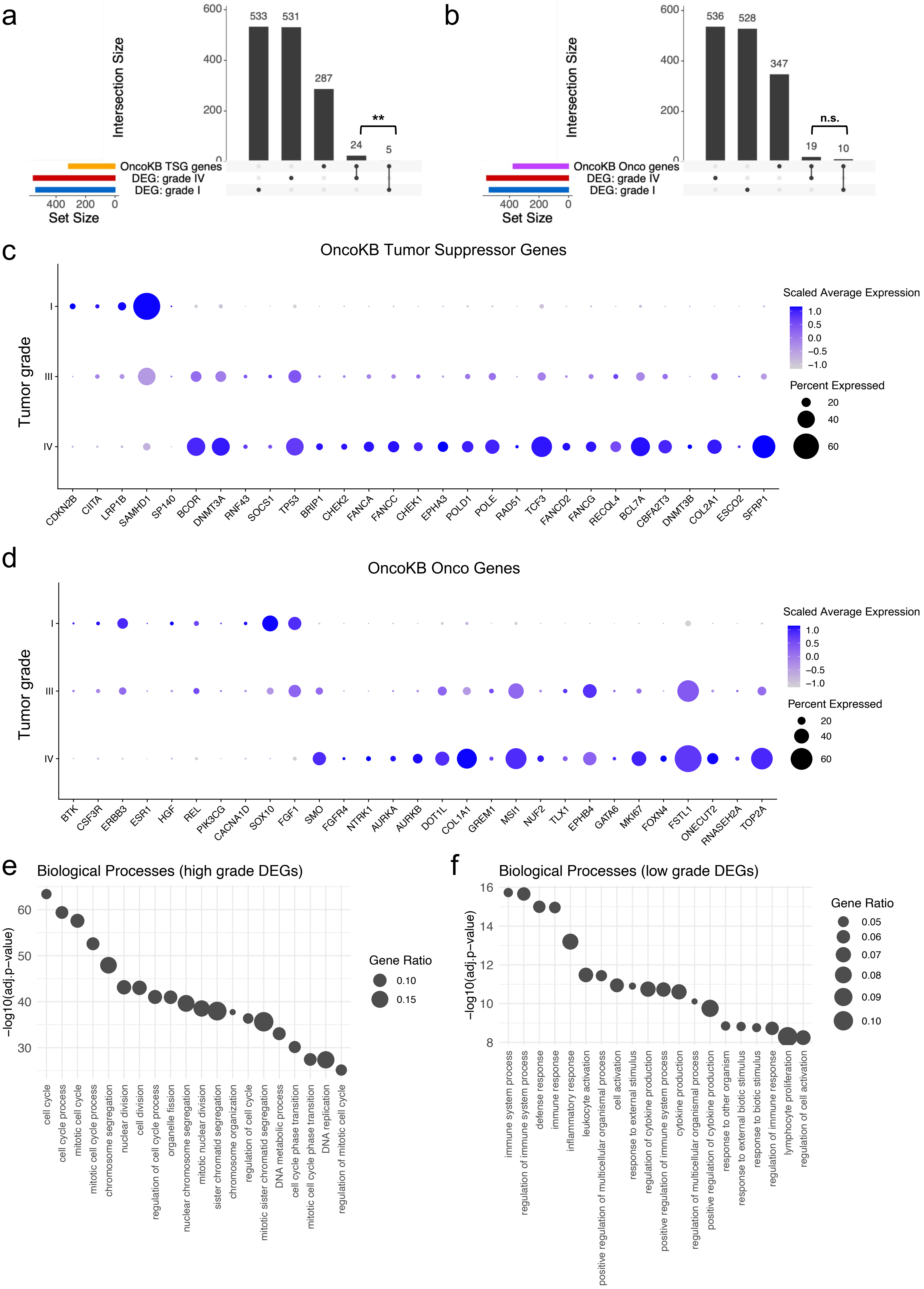

### Supplementary Figure 4

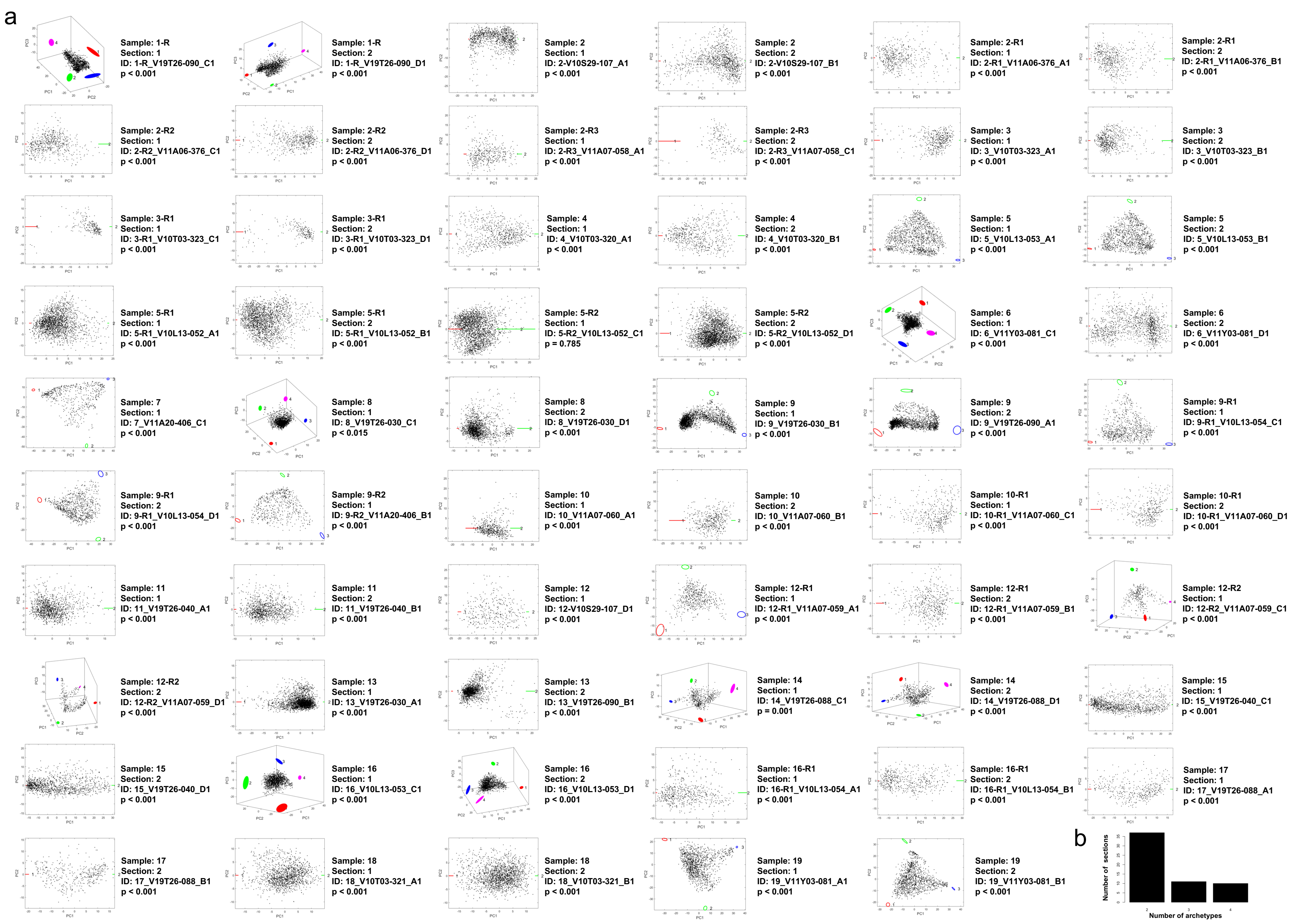

### Supplementary Figure 5

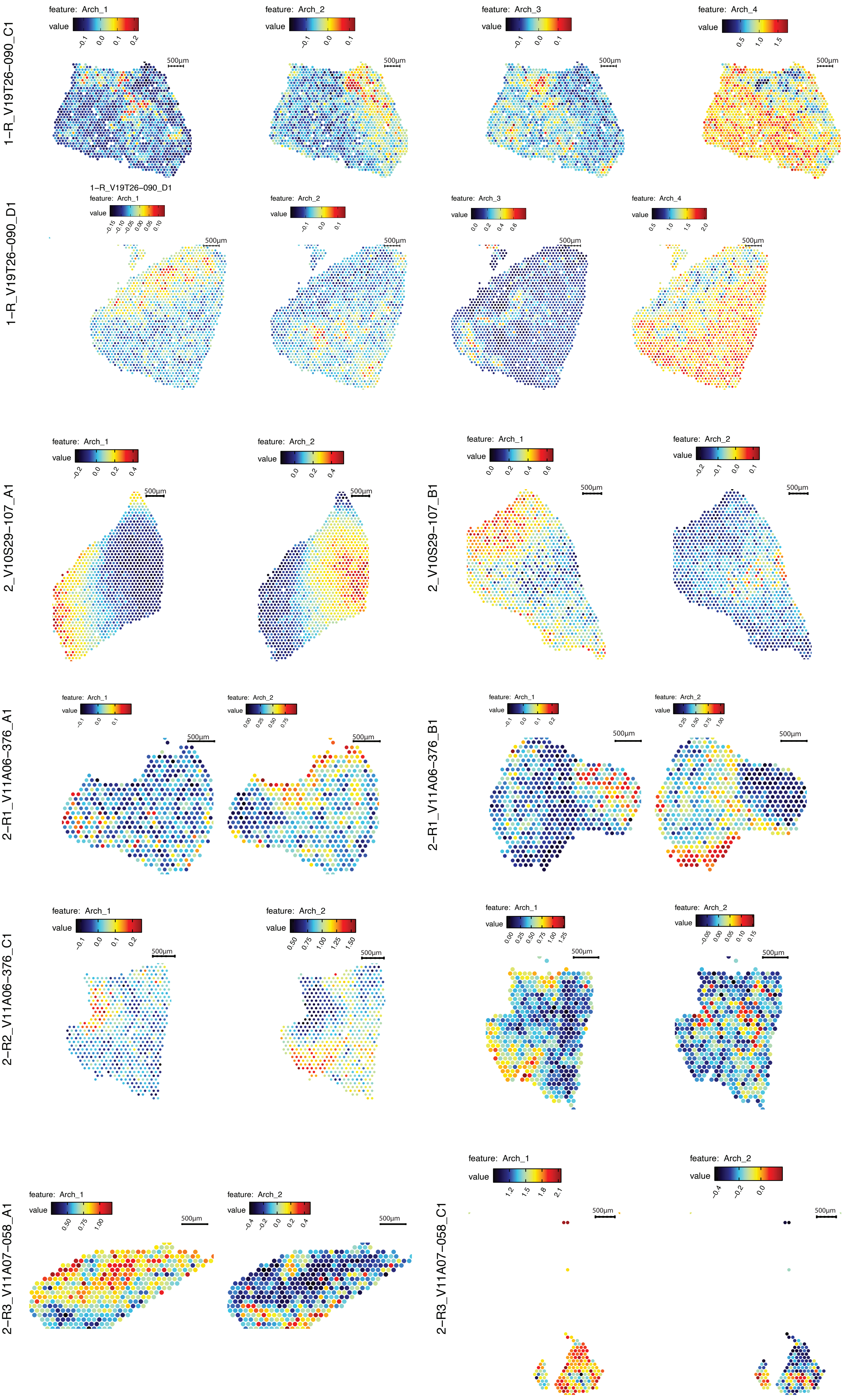

### Supplementary Figure 6

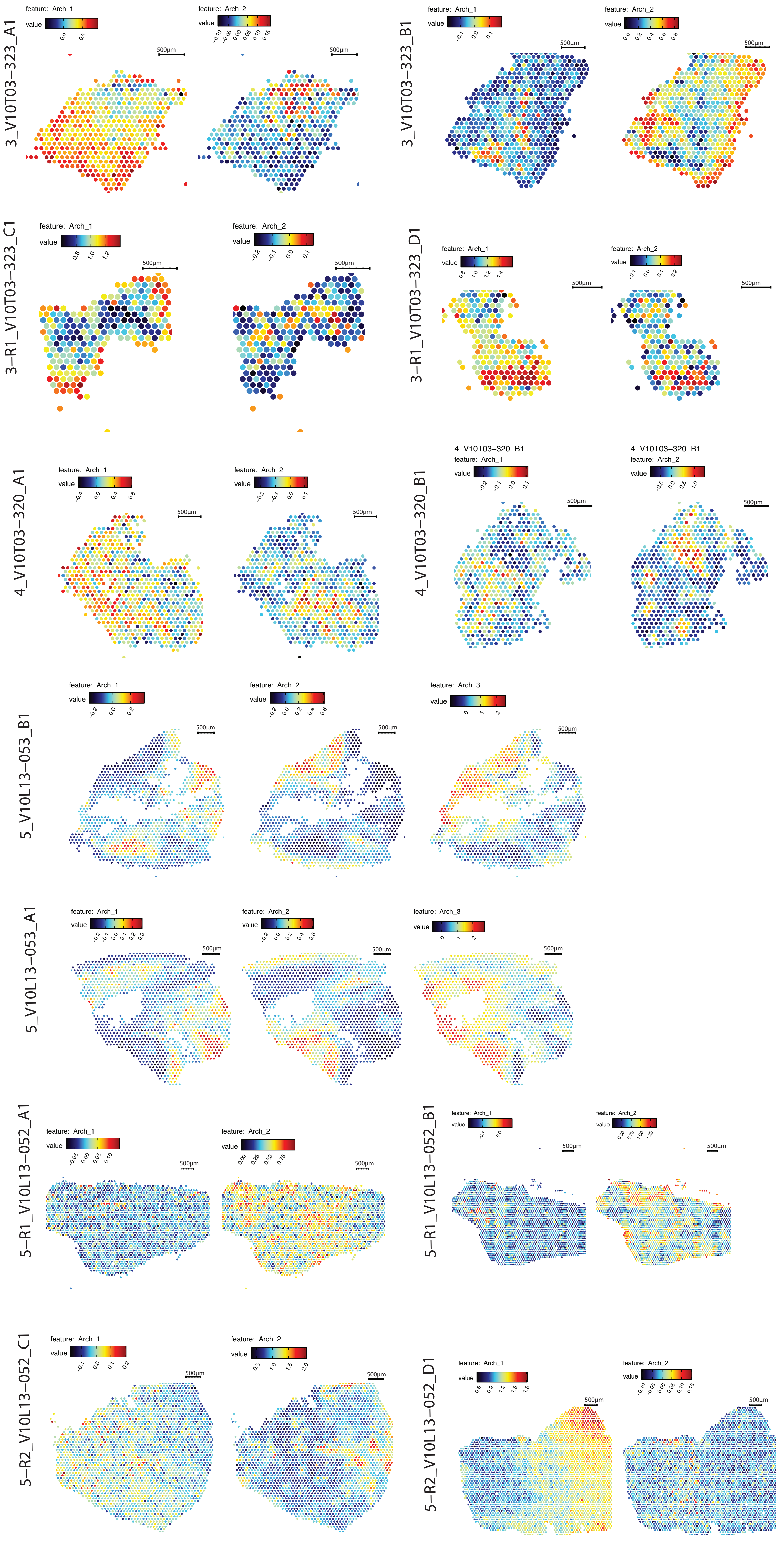

### Supplementary Figure 7

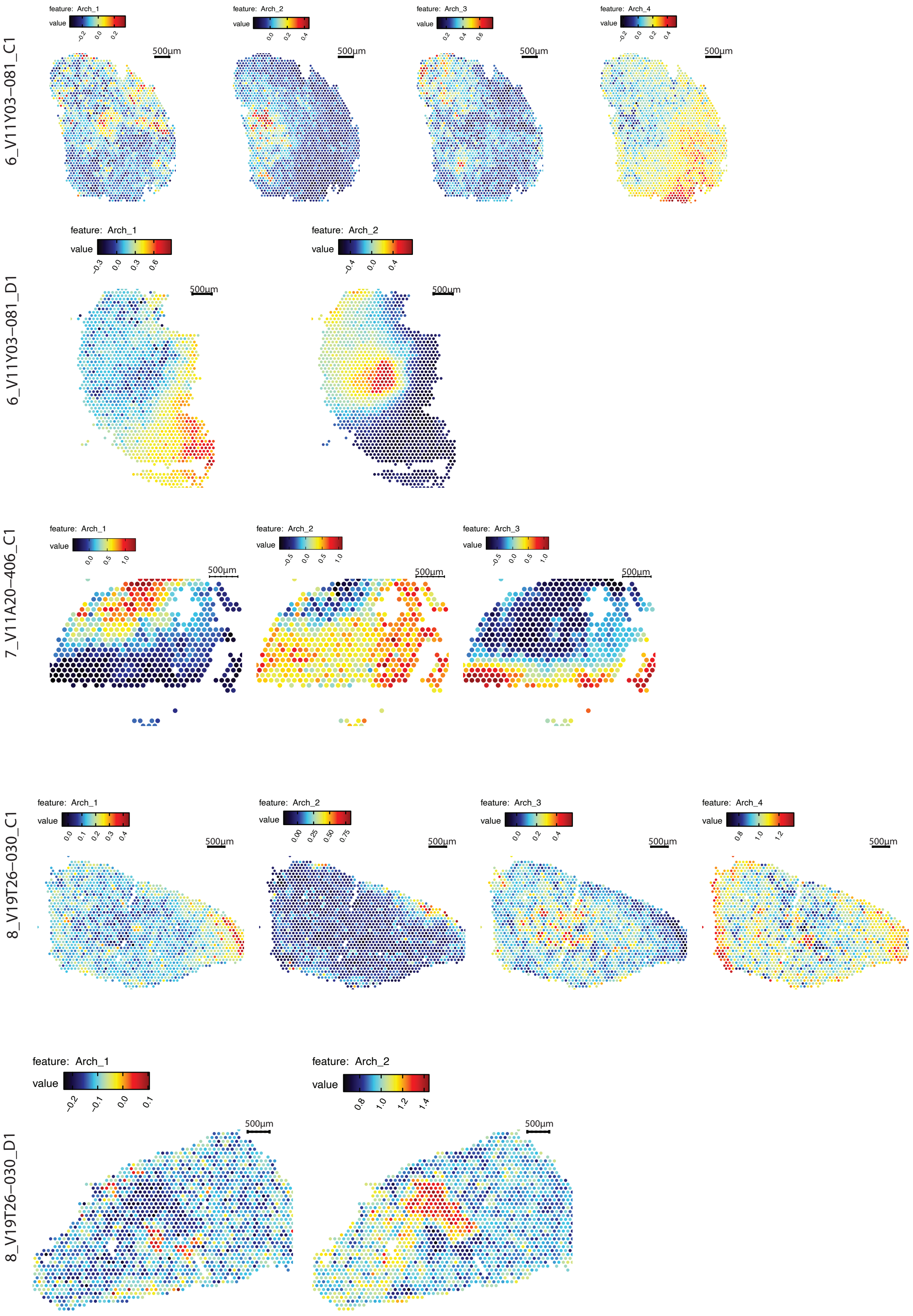

### Supplementary Figure 8

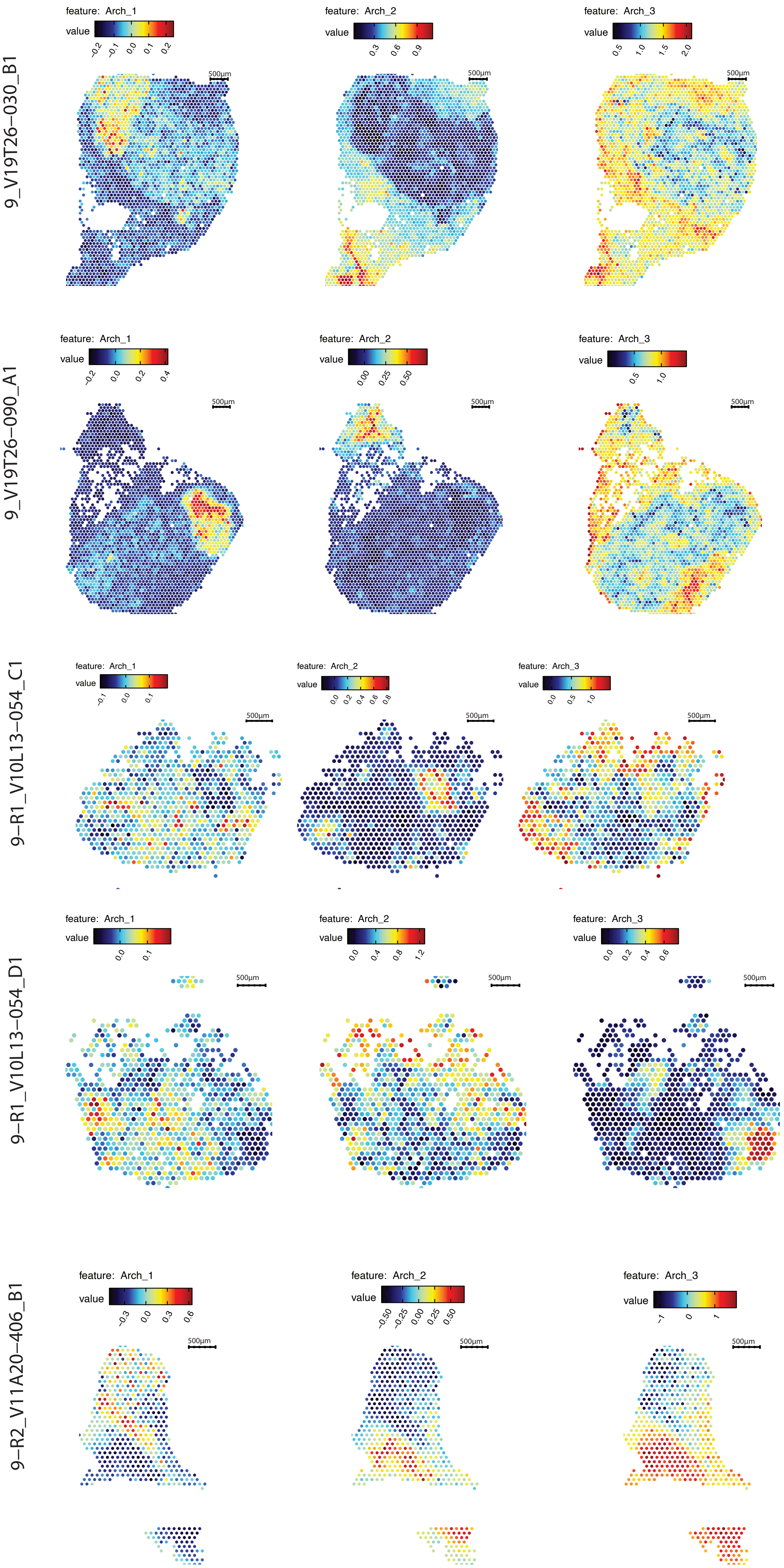

### Supplementary Figure 9

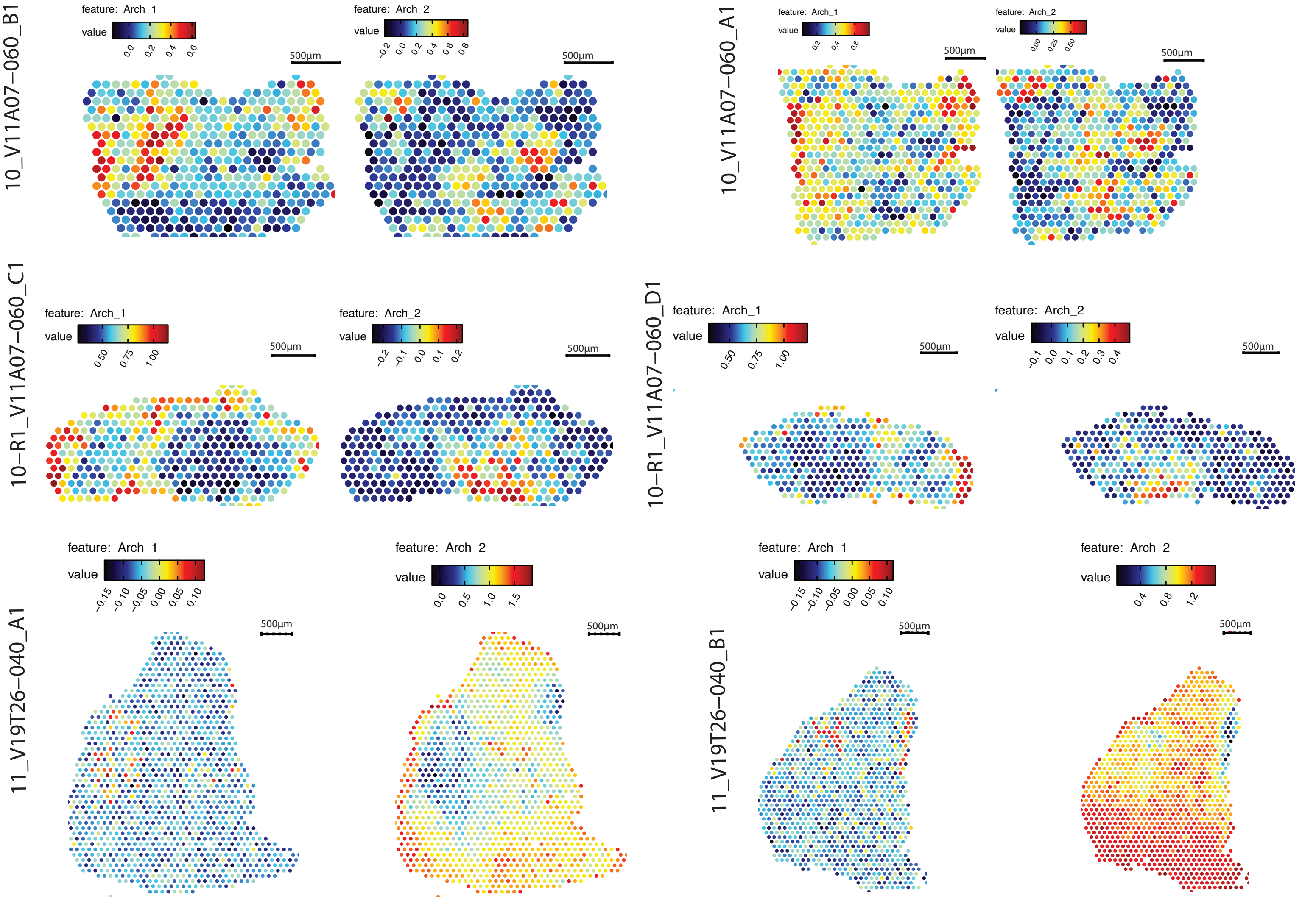

### Supplementary Figure 10

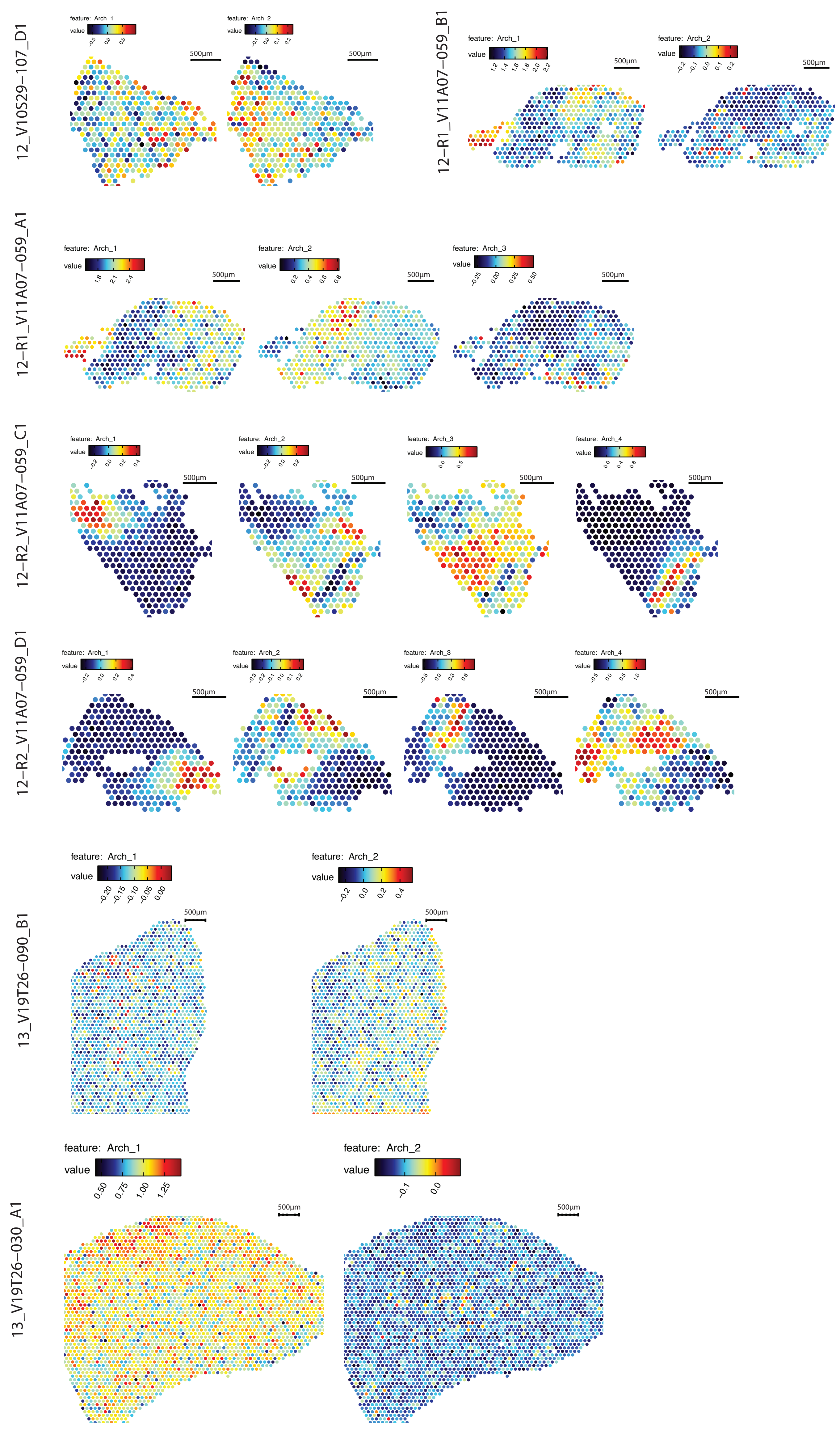

### Supplementary Figure 11

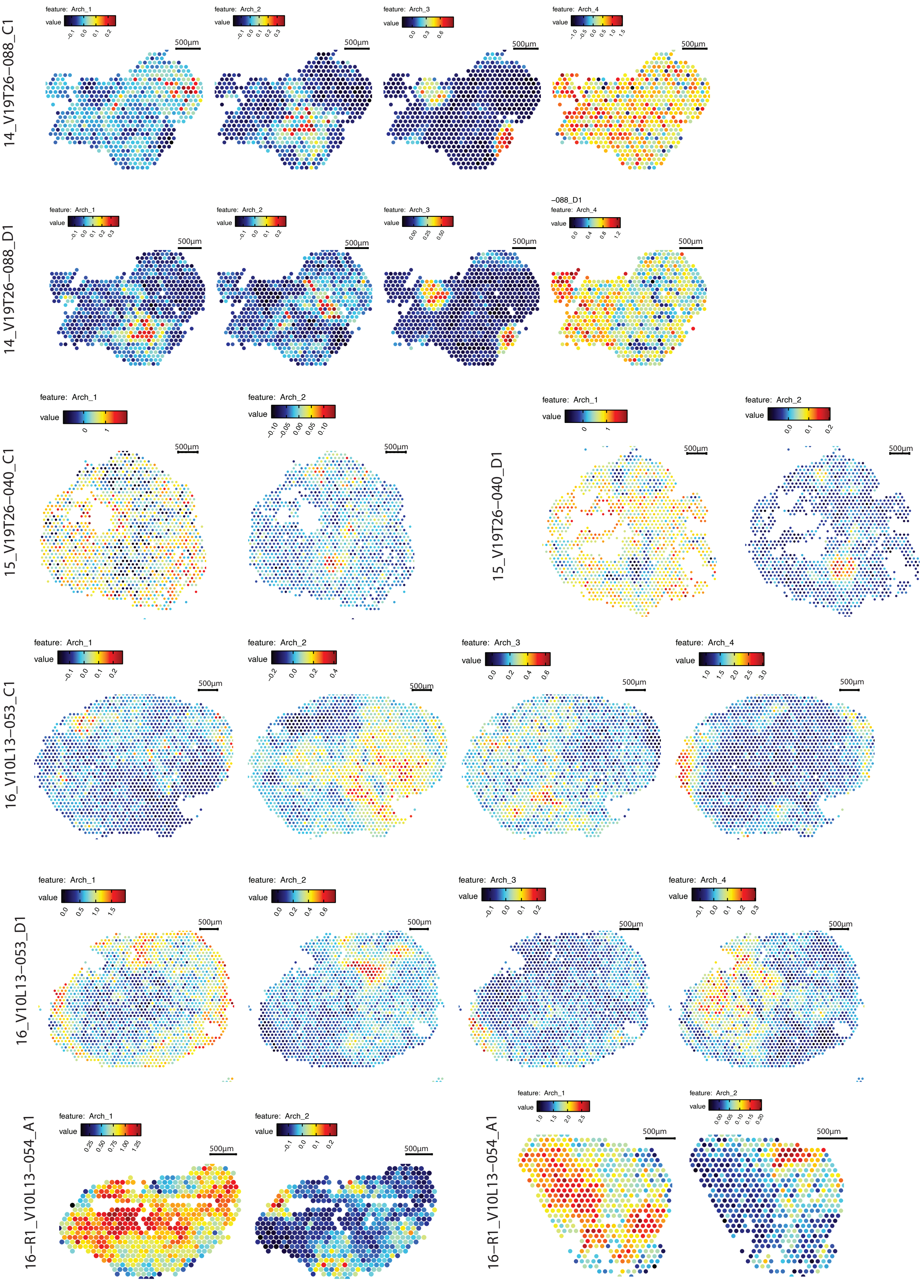

### Supplementary Figure 12

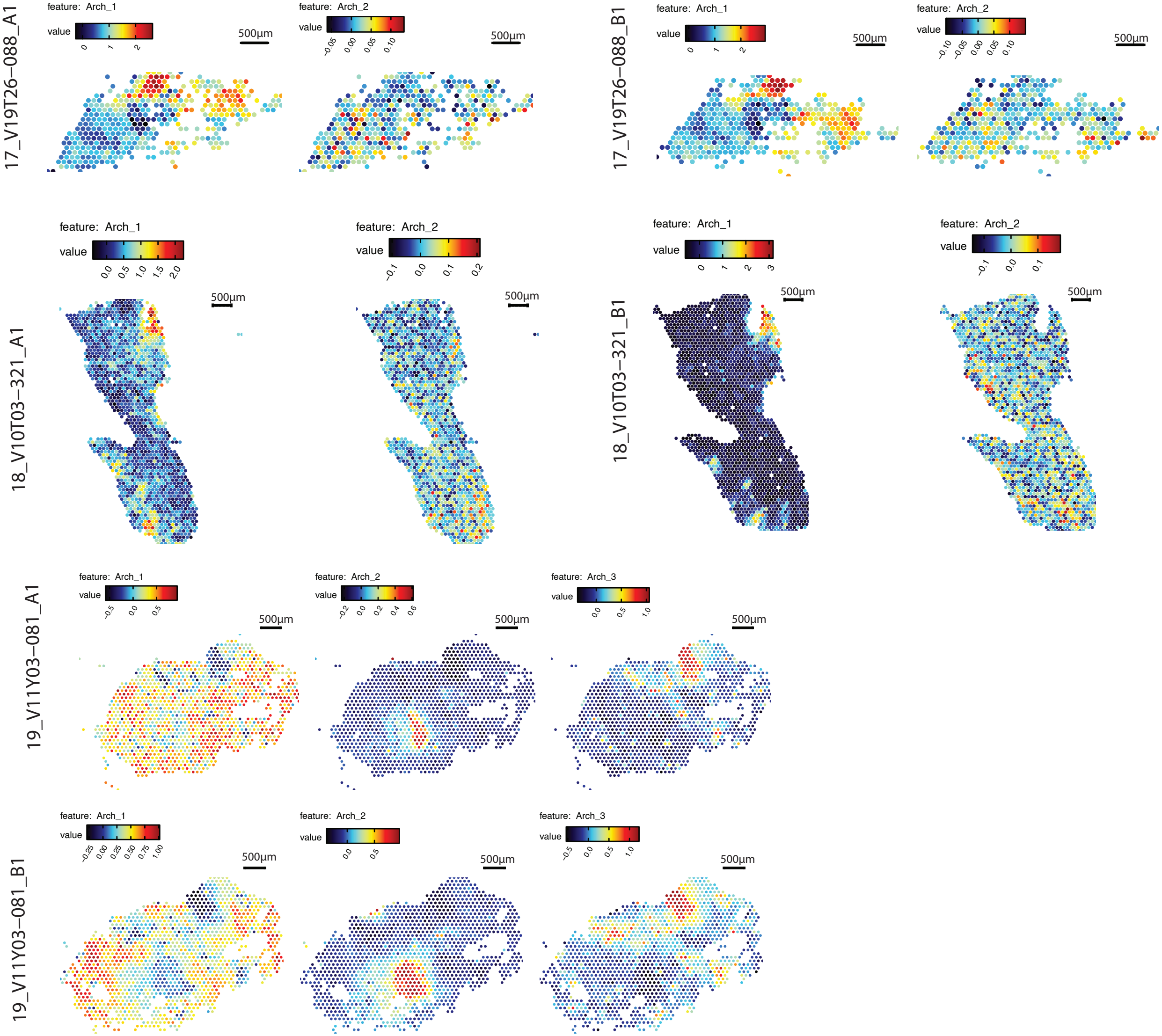

### Supplementary Figure 13

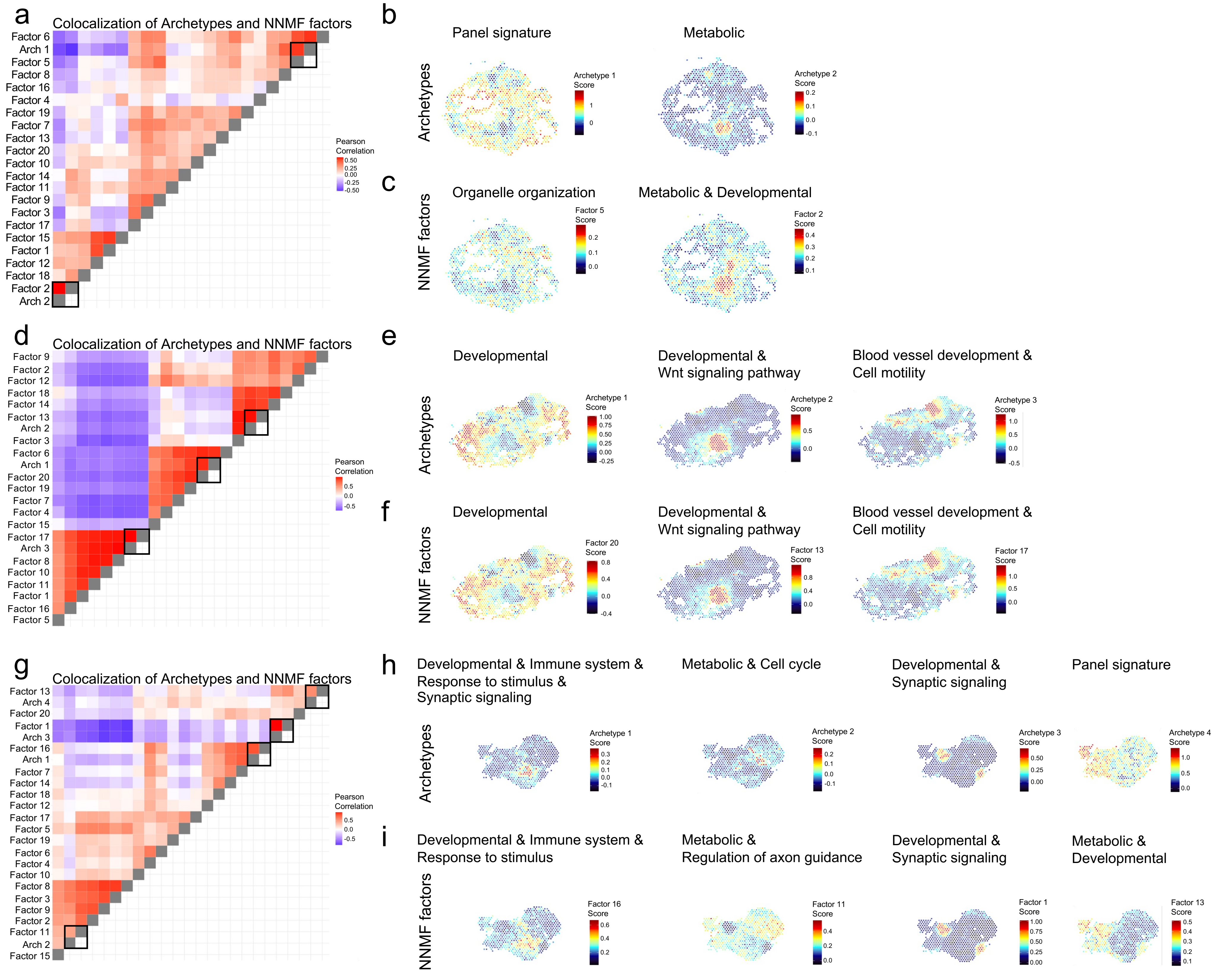

### Supplementary Figure 14

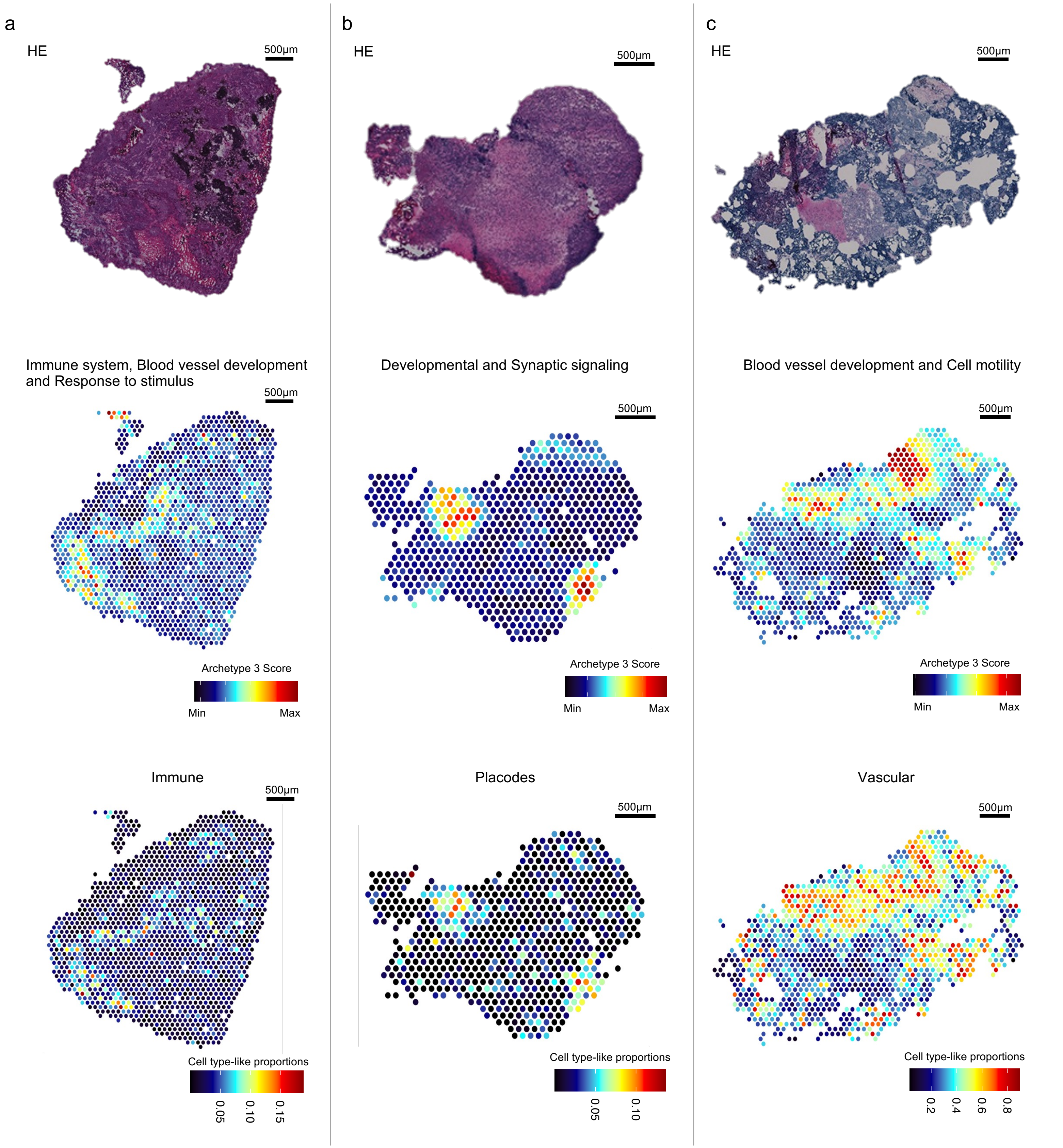

### Supplementary Figure 15

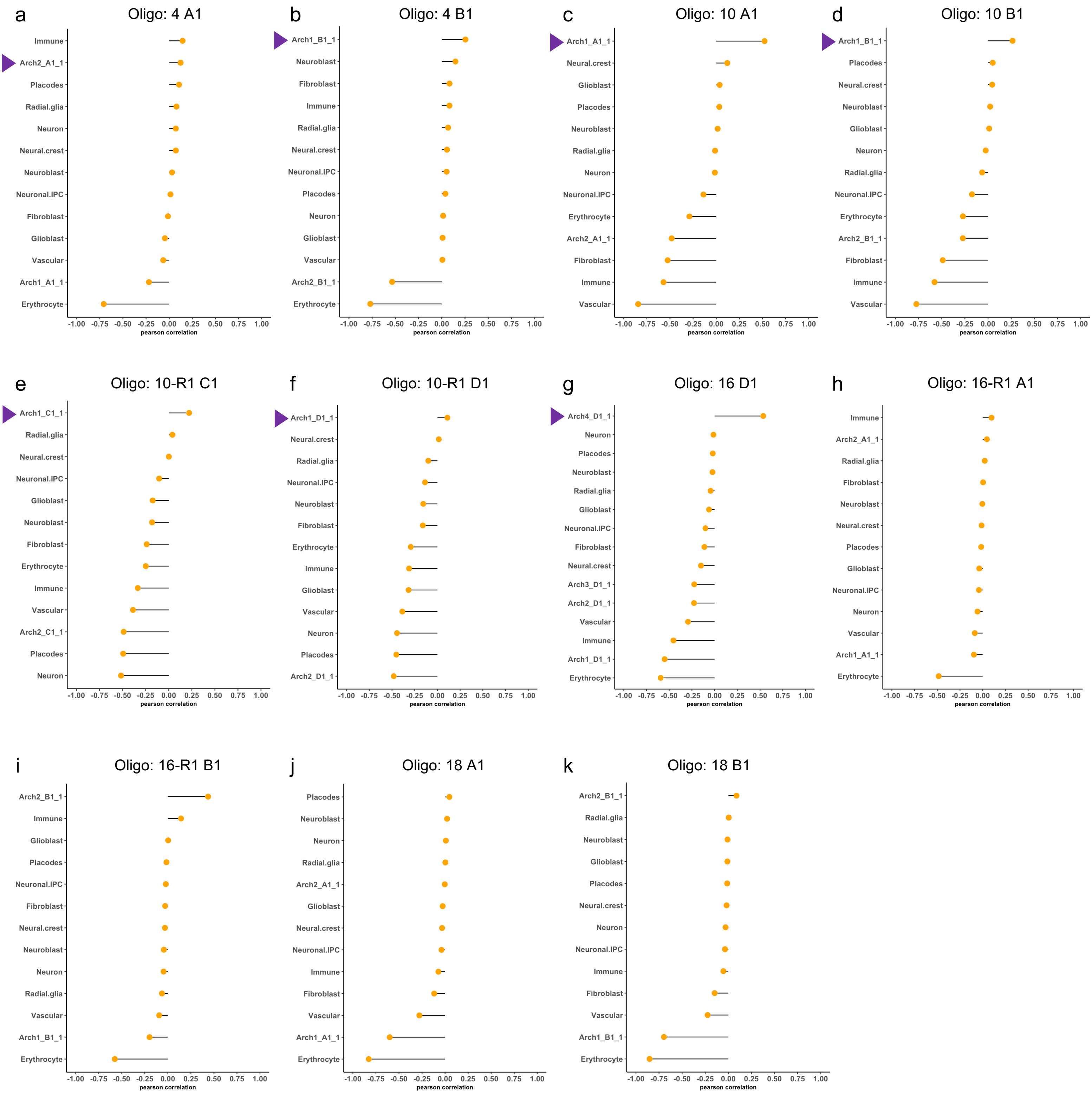

### Supplementary Figure 16

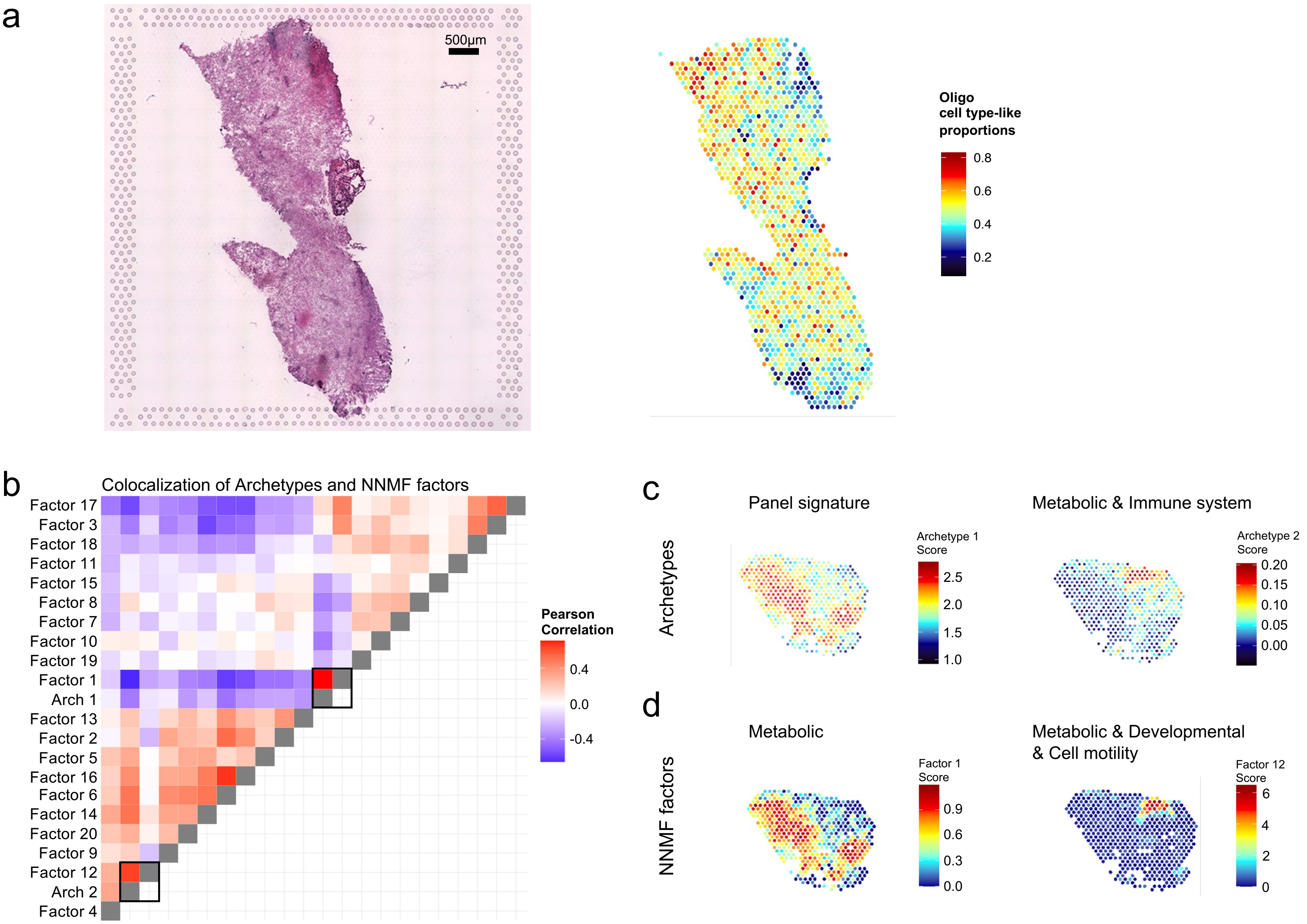

### Supplementary Figure 17

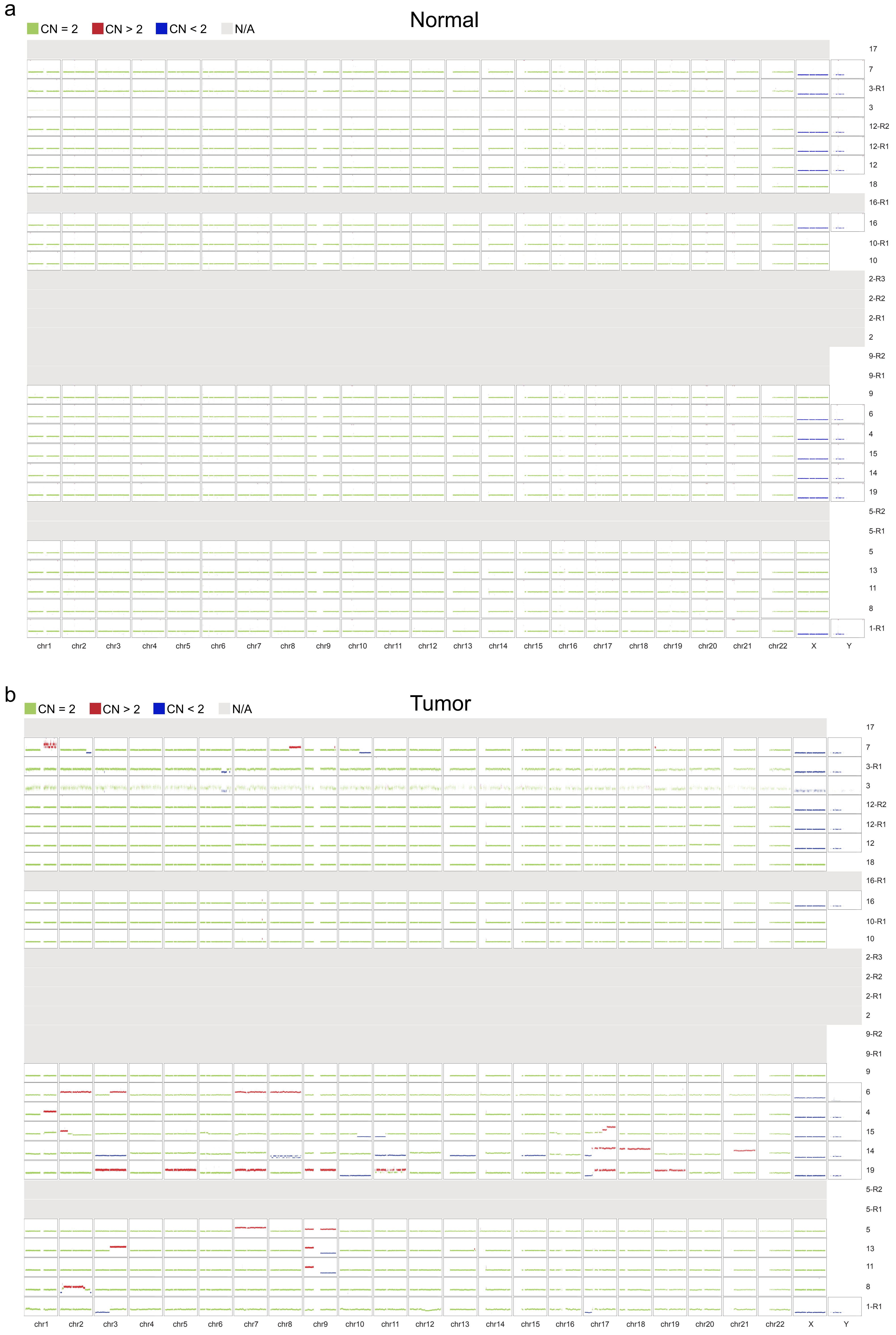

### Supplementary Figure 18

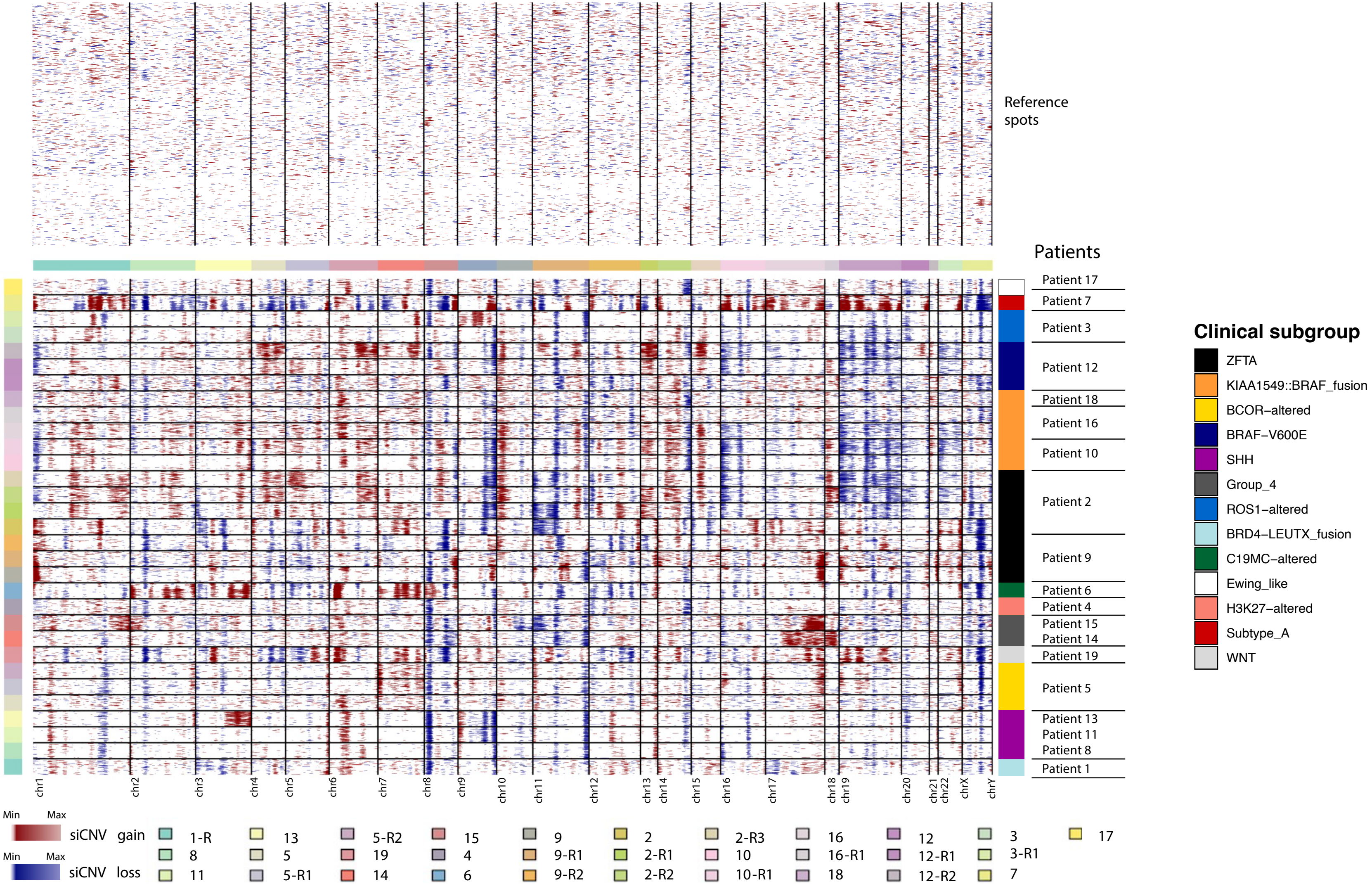

### Supplementary Figure 19

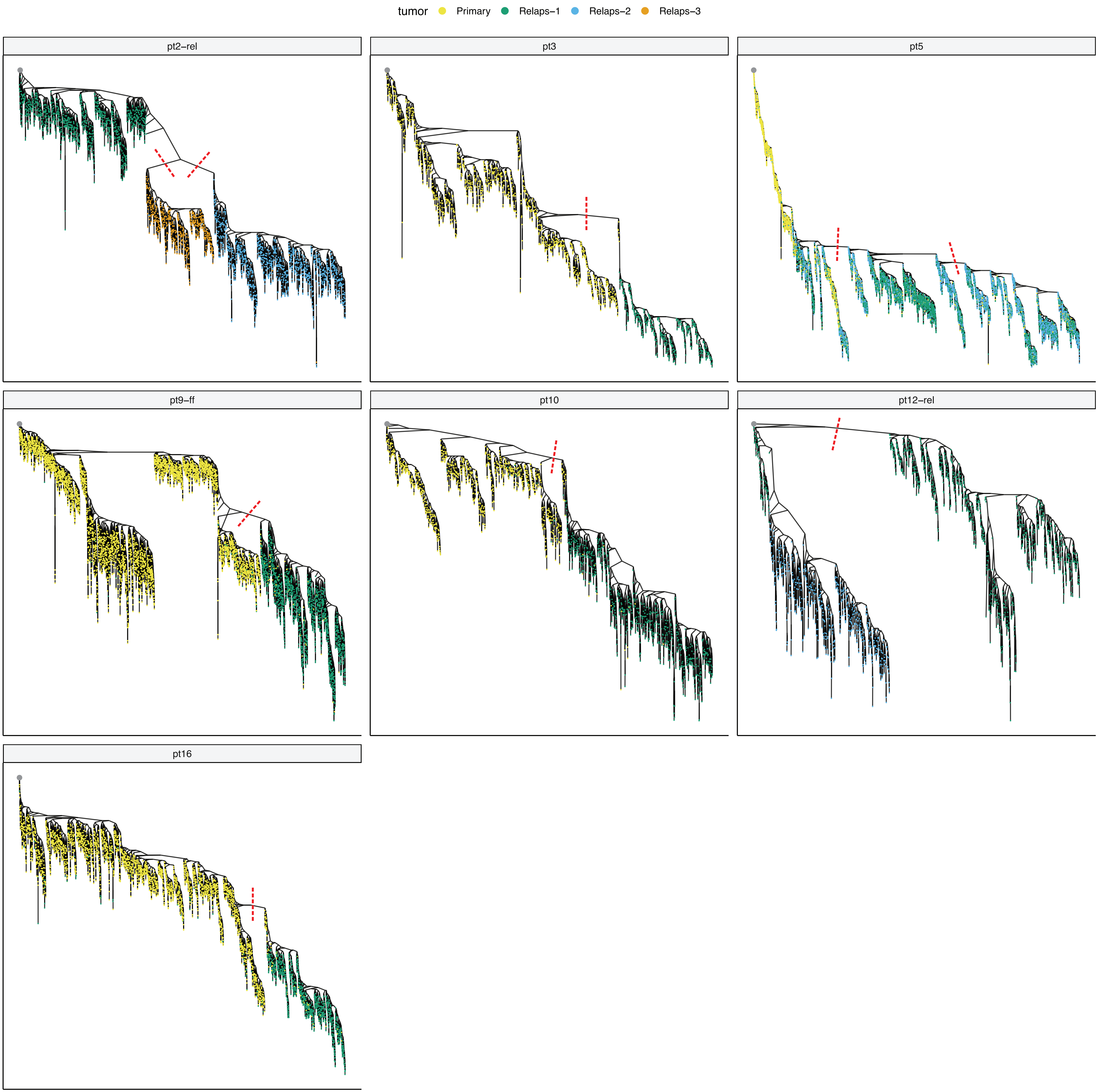

### Supplementary Figure 21

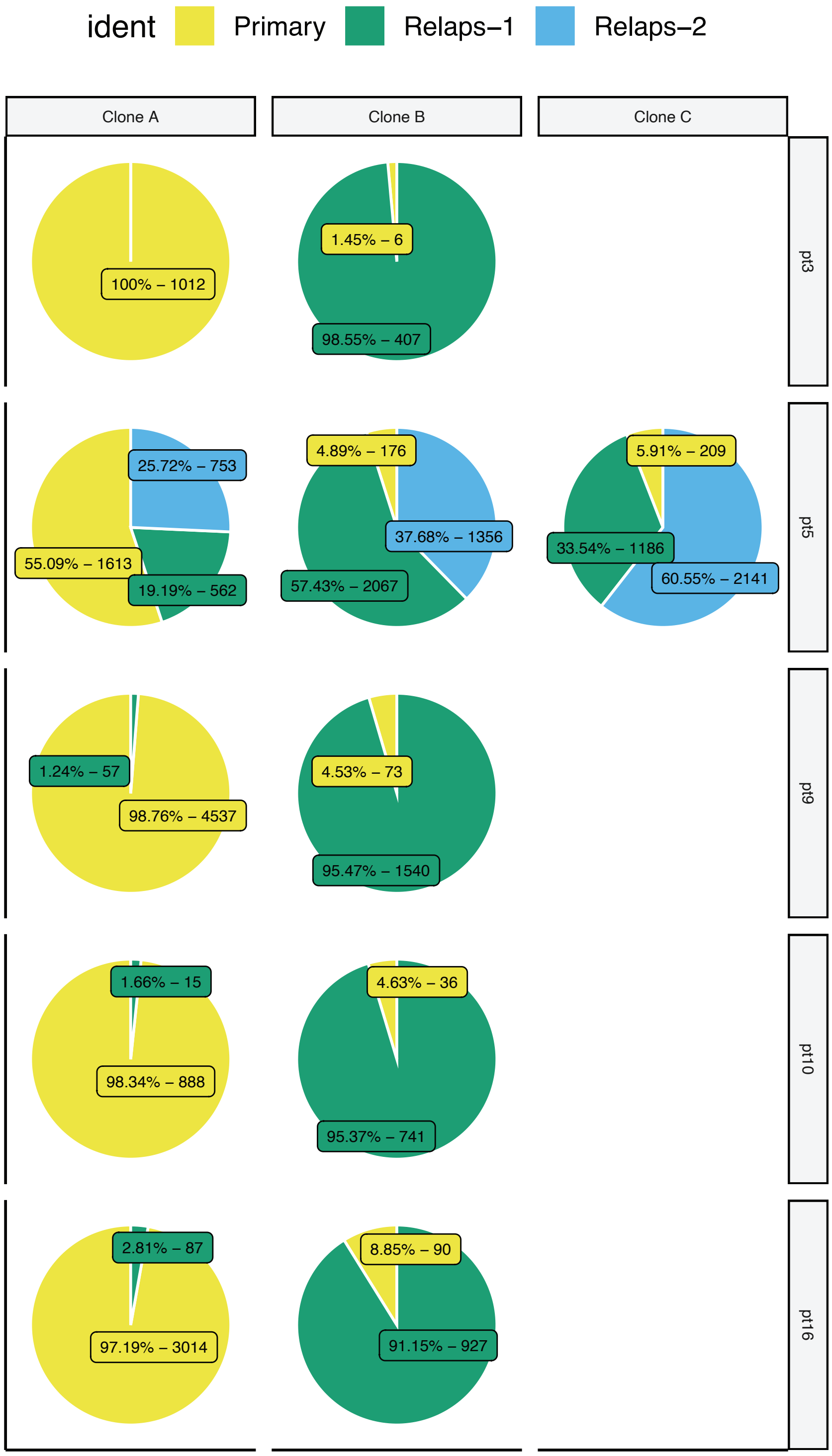
