## Supplementary Figure 20 for "Spatial mapping of pediatric brain tumors across diagnoses and relapses"

### patient 2 primaries

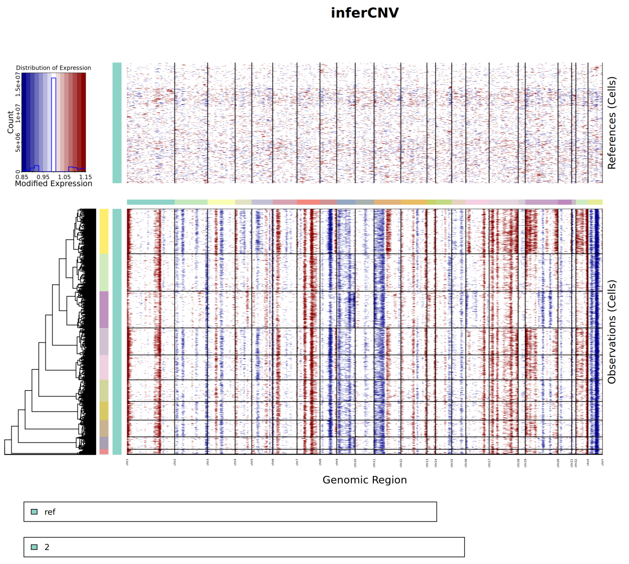


### patient 2 relapses

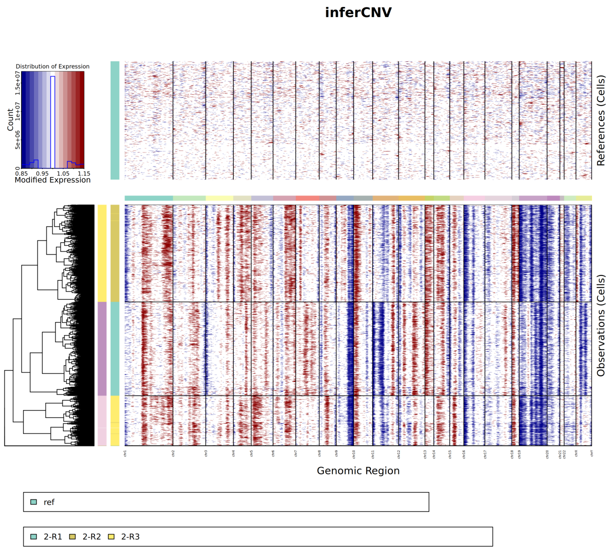


### patient 3

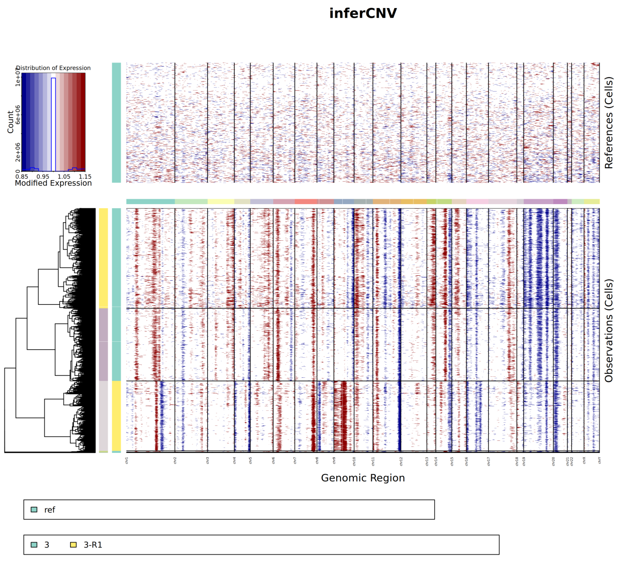


### patient 5

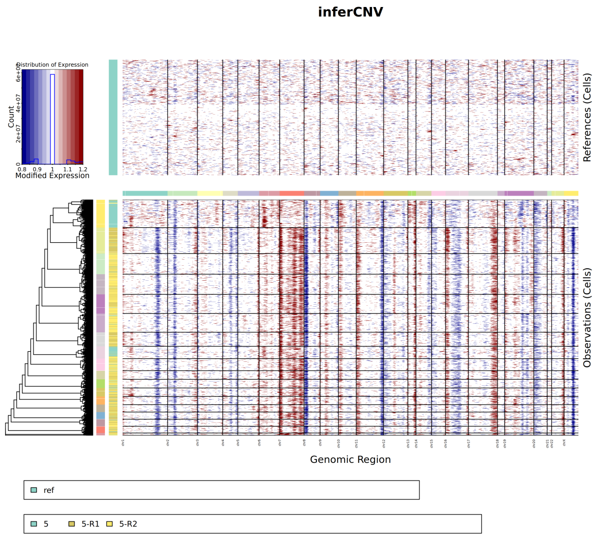


### patient 9 ff

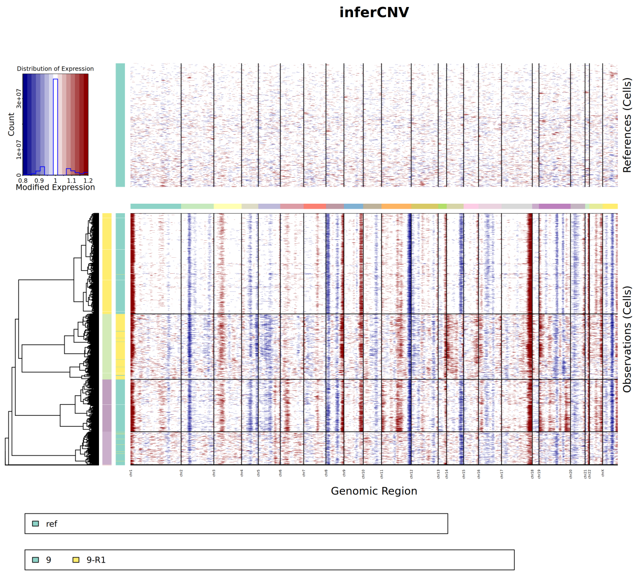


### patient 9 ffpe

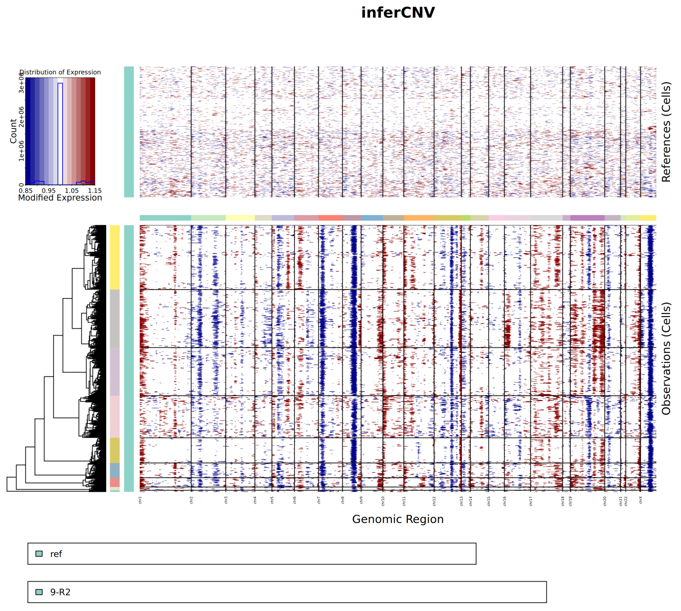


### patient 10

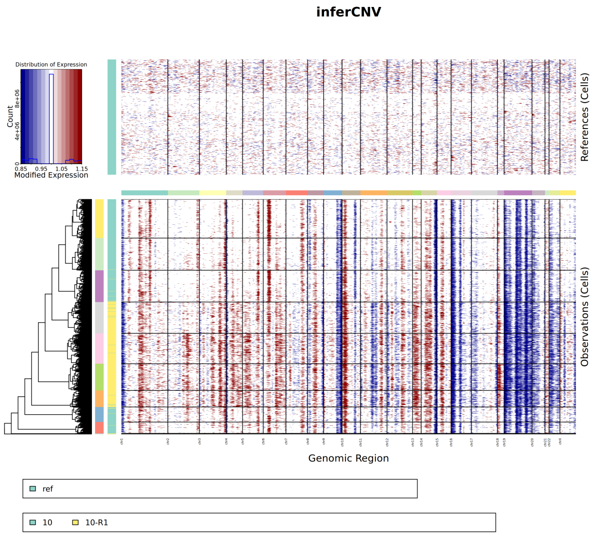


### patient 12 primaries

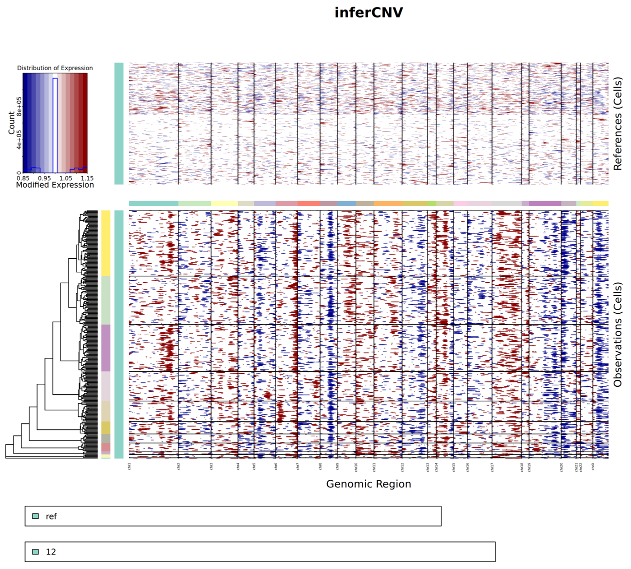


### patient 12 relapses

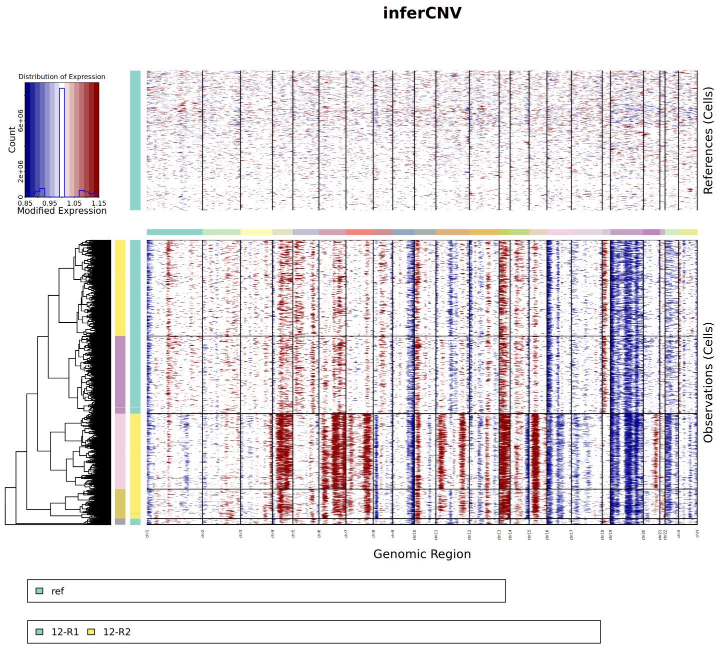


### patient 16

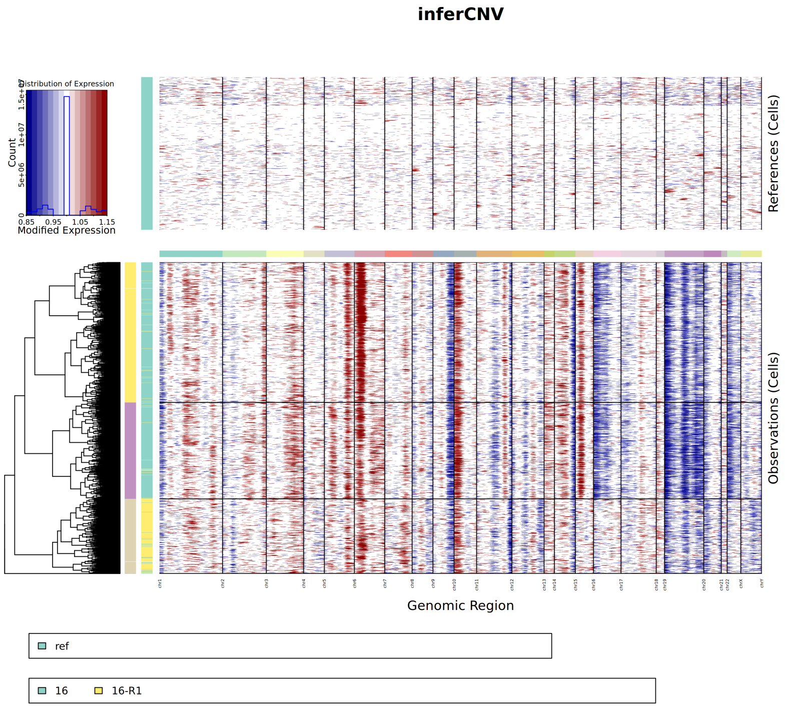
